# Glycan strand cleavage by a lytic transglycosylase, MltD contributes to the expansion of peptidoglycan in *Escherichia coli*

**DOI:** 10.1101/2023.05.04.539382

**Authors:** Moneca Kaul, Suraj Kumar Meher, Manjula Reddy

## Abstract

Peptidoglycan (PG) is a protective sac-like exoskeleton present in most bacterial cell walls. It is a large, covalently cross-linked mesh-like polymer made up of several glycan strands cross-bridged to each other by short peptide chains. Because PG forms a continuous mesh around the bacterial cytoplasmic membrane, opening the mesh is critical to generate space for the incorporation of new material during its expansion. In *Escherichia coli*, the ‘space-making activity’ is known to be achieved by cleavage of cross-links between the glycan strands by a set of redundant PG endopeptidases whose absence leads to rapid lysis and cell death. Here, we demonstrate a hitherto unknown role of glycan strand cleavage in making space for cell wall expansion in *E. coli*. We find that overexpression of a membrane-bound lytic transglycosylase, MltD that cuts the glycan polymers of the PG sacculus rescues the cell lysis caused by the absence of essential cross-link specific endopeptidases, MepS, MepM and MepH. Further detailed genetic and biochemical analysis revealed that MltD works in conjunction with cross-link specific endopeptidases to expand the PG sacculus. Interestingly, we find that cellular MltD levels are stringently controlled by two independent regulatory pathways. MltD undergoes regulated proteolysis by NlpI-Prc, a periplasmic adaptor-protease complex that specifically degrades two of the elongation-specific endopeptidases, MepS and MepH. In addition, MltD levels are post-transcriptionally controlled by RpoS, a stationary-phase specific sigma factor. Overall, our results show that coordinated cleavage of the glycan strands and the peptide cross-bridges facilitates the opening of the PG mesh for successful expansion of the cell wall during growth of a bacterium.

**AUTHOR SUMMARY:** Most bacteria are protected by a cell wall made up of peptidoglycan (PG), a mesh-like large polymer. PG consists of several linear glycan strands interlinked through short peptide chains to form a continuous meshwork around the bacterial cytoplasmic membrane. Because PG tightly encases the cytoplasmic membrane, the growth of a bacterial cell is coupled to the expansion of PG requiring the activity of hydrolytic enzymes that cleave PG cross-links to make space for incorporation of new PG material. ln *E. coli*, a set of redundant cross-link specific endopeptidases are known to be crucial for expansion of PG. In this study, we show that cleavage of the glycan polymers by MltD, a glycan cleaving enzyme compensates the absence of cross-link cleavage and contributes to the expansion of PG. Overall, our work shows a previously unknown role of glycan hydrolases in cell wall expansion identifying these as potential targets for development of cell wall-specific antimicrobial agents.

## INTRODUCTION

Most bacteria are surrounded by a protective exoskeleton, the peptidoglycan (PG or murein) that protects cells from lysis by internal osmotic pressure and harsh environmental conditions. In Gram-negative bacteria, PG is located in the periplasmic space in between the inner membrane (IM) that encloses the cytoplasm and a surface-exposed outer membrane (OM). PG is a covalently closed, mesh-like macromolecule that closely encases the IM providing shape to the bacterial cells. Structurally, it is made up of several linear glycan polymers containing repeating disaccharide units of N-acetylglucosamine (GlcNAc) and N-acetylmuramic acid (MurNAc) residues covalently bonded by β-1,4 glycosidic linkage. Each MurNAc residue of the glycans is attached to a short peptide chain of three to five amino acids with a pentapeptide consisting of, L-alanine^1^−D-glutamate^2^−meso-diaminopimelic acid (mDAP)^3^−D-alanine (D-ala)^4^−D-ala^5^. Approximately 40% of the peptide chains in the PG are cross-linked to each other; of these, approximately 90-93% form between the D-ala^4^ of one peptide chain and the mDAP^3^ of the adjacent chain (D-ala−mDAP or 4,3 cross-links) by the catalytic activity of either class A (PBP1A, PBP1B) or class B (PBP2, PBP3) PG synthases whereas a small fraction (7-10%) form between the two mDAP residues (mDAP−mDAP or 3,3 cross-linking) by the activity of L,D-transpeptidases, LdtD and LdtE [1–4].

Because the PG forms a continuous network around the IM, the growth and enlargement of a bacterial cell is tightly coupled to the expansion of PG sacculus [5]. In several Gram-negative bacteria, cleavage of cross-links by PG hydrolytic enzymes has been shown to be essential for their growth and viability suggesting that the hydrolysis of cross-links is crucial to make space for the incorporation of nascent glycan strands during PG expansion [6–8]. *E. coli* encodes several 4,3 cross-link specific PG hydrolases (termed D,D-endopeptidases) which include, MepA (*mepA*), MepS (*mepS*), MepM (*mepM*), MepH (*mepH*), Pbp4 (*dacB*), Pbp7 (*pbpG*) and AmpH (*ampH*) [6,9,10]. Among these, absence of MepS, -M and -H leads to rapid lysis and cell death in *E. coli* demonstrating their essentiality at normal physiological growth conditions [6,11].

Although, cross-link hydrolysis is crucial for the expansion of PG, it needs to be controlled stringently as unfettered cleavage leads to lethal degradation and rupture of the PG sacculus. We previously showed that a key elongation-specific endopeptidase, MepS in *E. coli* is controlled at the step of post-translational stability by a periplasmic proteolytic system comprising an OM lipoprotein, NlpI and a soluble C-terminal specific protease, Prc [12]. Here, NlpI serves as an adaptor protein to bind both MepS and Prc to bring them into proximity to facilitate degradation of MepS by Prc [12,13]. The NlpI-Prc system is also conserved in *Pseudomonas aeruginosa* in which the homologs of MepS and MepM are cleaved by the protease CtpA through a lipoprotein adaptor LbcA [8,14,15].

In addition to endopeptidases, *E. coli* encodes several classes of PG hydrolases including carboxypeptidases, lytic transglycosylases and amidases which function in PG maturation, remodelling, turnover and daughter cell separation [9,10,16]. The lytic transglycosylases or LTs catalyse the nonhydrolytic cleavage of β,1-4 glycosidic linkages between GlcNAc and MurNAc residues of PG with concomitant formation of a cyclic1,6-anhydro ring at the MurNAc residue [16–18]. Due to this distinct catalytic activity of LTs, the termini of most glycan strands in *E. coli* contain 1,6-anhydro MurNAc residues. LTs are believed to play a predominant role in PG recycling as the 1,6 anhydro muropeptides generated by their activity are transported through AmpG, an IM-permease to be further utilised for the synthesis of PG precursors in the cytosol [19]. Based on the substrate specificity, LTs are either endolytic (cleaving within the glycan strand) or exolytic (cleaving from the ends of a glycan chain) [16,17]. *E. coli* encodes eight lytic transglycosylases, of which MltA, -B, -C, -D, -E, -F are OM-associated; Slt is soluble periplasmic; and MltG is IM-anchored [20]. In addition, a division-associated glycosyl hydrolase, DigH which specifically targets denuded glycan strands at the septum is known [21]. It is not yet clear why *E. coli* has such a large repertoire of glycan hydrolases; however, recent evidence suggests that they have distinct substrate-specificity and function in discrete PG pathways [20–24].

In this study, we show that glycan strand cleavage by the lytic transglycosylase, MltD [25] contributes to the PG expansion in *E. coli*. During an attempt to understand the mechanism of PG enlargement, we observed that overexpression of MltD, an OM lipoprotein compensates the absence of cross-link cleaving endopeptidases, MepS, MepM and MepH. Further analysis revealed that MltD plays a role in optimal growth and enlargement of PG sacculus along with the cross-link specific endopeptidases. Interestingly, we find that cellular MltD levels are maintained by two parallel regulatory pathways. It is controlled at the step of post-translational stability by NlpI-Prc mediated proteolysis and in addition, the MltD abundance in the stationary-phase cells is governed by the stationary-phase specific sigma factor, RpoS. Biochemical analysis shows MltD is an endolytic LT that is rapidly degraded by NlpI-Prc in vitro. Overall, our results show that MltD, a stringently controlled lytic transglycosylase, contributes to PG enlargement in conjunction with the cross-link specific endopeptidases to effectively make space for the insertion of nascent glycan material during PG expansion.

## RESULTS

### Absence of NlpI-Prc proteolytic machinery compensates the endopeptidase insufficiency in E. coli

Absence of two major cross-link specific PG endopeptidases, MepS and MepM leads to rapid lysis of *E. coli* cells growing in rich media such as LB [6,11]. To understand the basis of cell lysis in these mutants, we sought to identify suppressor mutations that conferred viability to *mepS mepM* double deletion mutants on LB medium. Approximately 30 spontaneous suppressor mutants from two independently grown cultures were isolated and the mutations were identified using conventional gene mapping techniques (as described in Materials and Methods). Surprisingly, all these mutations were found to be recessive alleles of either *nlpI* or *prc* (Fig. 1A). In support of this observation, the deletion alleles of *nlpI* or *prc* also suppressed the growth defects of *mepS mepM* double mutants on LB (Fig. 1A). NlpI and Prc form a proteolytic complex in the periplasm in which NlpI functions as an adaptor protein to bring its substrate MepS to the proximity of Prc for its degradation [12,13]. The rescue of *mepS mepM* mutant by *nlpI* or *prc* gene deletions suggested the existence of an alternate PG endopeptidase that is stabilized in the absence of NlpI-Prc proteolytic machinery and thereby compensating the loss of MepS and MepM.

**Figure 1.**
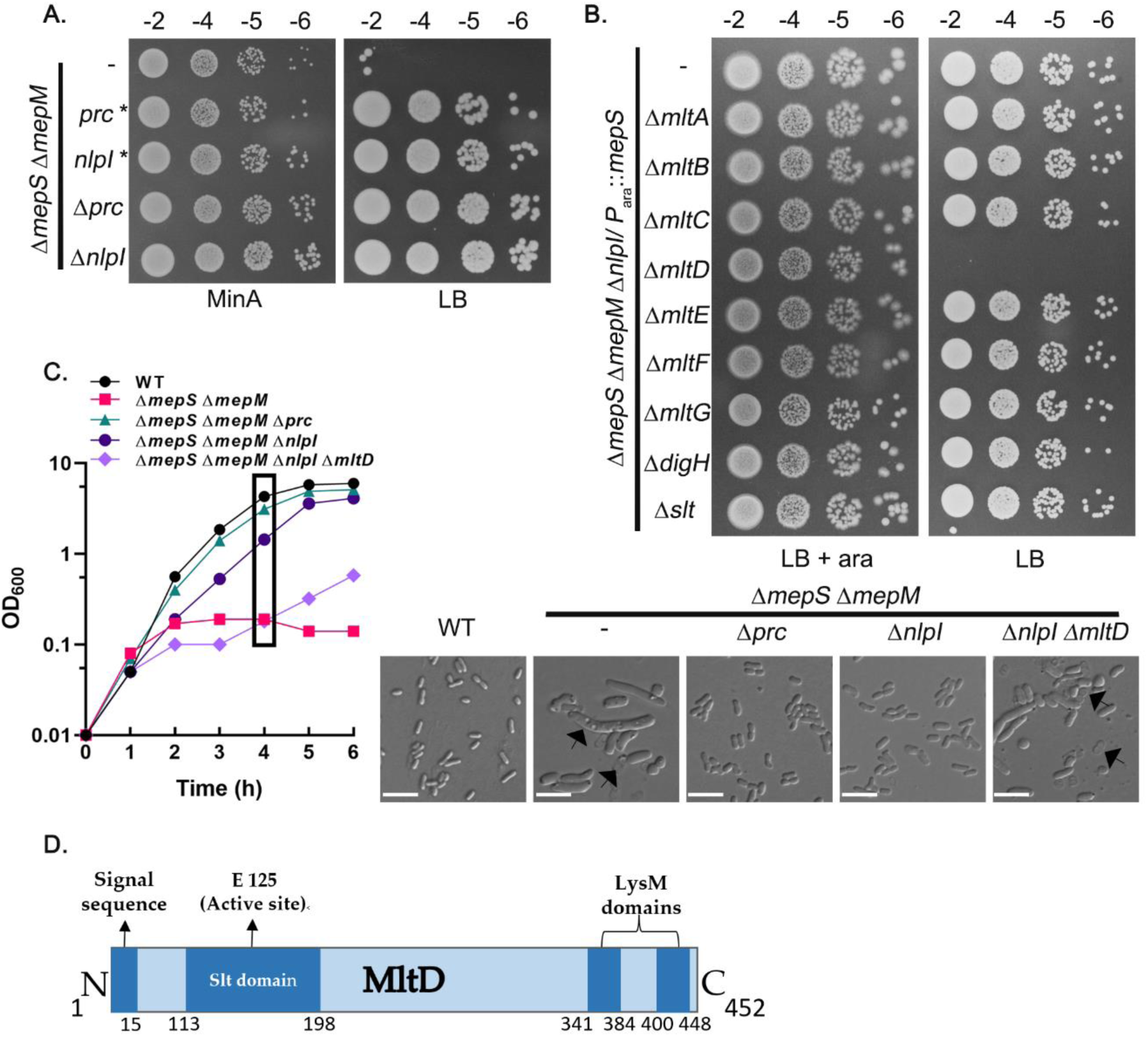
Absence of NlpI-Prc system rescues the growth defects of *mepS mepM* mutant through MltD. (A) *mepS mepM* double mutant and its derivatives lacking either *nlpI* or *prc* were tested for viability on Min-A and LB plates at 37°C. Cells were grown overnight in Min-A, serially diluted and 4 µl of each dilution was spotted on indicated plates and grown overnight. (B) Δ*mepS* Δ*mepM* Δ*nlpI*/ P_ara::_*mepS* and its derivatives carrying deletion of each of the LTs were tested for viability on LB with and without arabinose (0.2%) at 37°C. (C) Cultures of WT and its mutant derivatives were grown overnight in Min-A. Next day, they were washed and diluted 1:250 into fresh LB and growth was monitored without arabinose. Cells were collected after 4 h growth (as indicated in the box) and visualized by DIC microscopy. Arrows indicate cell lysis. Scale bar equals 5 µm. (D) Depiction of structural features of MltD protein indicating its transglycosylase domain and two C-terminal LysM repeats.

To identify the factor(s) responsible for the suppression, we deleted each of the known PG endopeptidases of *E. coli* (MepA, MepH, MepK, PBP4, PBP7 or AmpH) from *mepS mepM nlpI* triple deletion mutants; however, none of these deletions were able to abolish the growth of this triple mutant suggesting that the stabilization of these alternate endopeptidases is not the basis of the suppression (Fig. S1A). In contrast to a recent observation [11] which showed that stabilization of MepH as the basis of suppression in Δ*mepS* Δ*mepM* double mutant lacking NlpI or Prc, our results show that absence of NlpI-Prc system is still able to suppress the growth defect of *mepS*, -*M* and -*H* triple mutants (Fig. S1B).

### Overexpression of a lytic transglycosylase, MltD compensates the absence of essential cross-link specific endopeptidases

To identify the factor(s) potentially regulated by NlpI-Prc system, we took a candidate approach and introduced deletion of the genes encoding several PG hydrolases each involved in the synthesis, recycling, or turnover of PG into the *mepS mepM nlpI* triple mutant. Of all the deletions tested, we observed that deletion of *mltD*, a gene encoding a membrane-bound murein lytic transglycosylase was able to totally abrogate the growth of the triple mutant on LB (Fig. 1B, 1C) suggesting that MltD could be the putative hydrolase responsible for the suppression of *mepS mepM* double mutant in the absence of NlpI. On the contrary, deletion of MltD in *mepS mepM prc* had only a minor effect on its viability suggesting Prc may have additional NlpI-independent substrates that function in PG expansion (Fig. S1C). However, deletion of any other LT encoded by *E. coli* did not affect the viability of *mepS mepM nlpI* triple mutant (Fig. 1B).

Next, we examined whether increasing the expression of any of the LTs can overcome the lethality of *mepS mepM* double mutant. For this purpose, we overexpressed each of the LT (cloned downstream of an IPTG-inducible promoter; ASKA collection) [26] in *mepS mepM* strain and examined their growth. Of all the plasmids tested, only the plasmid encoding MltD was able to restore viability to this double mutant (Fig. 2A; Fig. S1D). To confirm this observation further, we cloned *mltD* in a medium-copy vector, pTrc99a downstream of an IPTG-inducible *trc* promoter (P_trc_::). This clone also behaved like that of ASKA-*mltD* and conferred suppression to both *mepS* and *mepS mepM* double mutants (Fig. 2B, 2D). Notably, overexpression of MltD rescued the growth of *mepS mepM mepH* triple mutant signifying the ability of MltD to compensate the lack of all the crucial cross-link cleaving endopeptidases of *E. coli* (Fig. 2C). To examine whether the enzymatic activity of MltD is obligatory for the suppression, we made catalytically inactive mutants of *mltD* (E125A and E125K; Fig. 1D) and find that these inactive variants do not confer growth advantage showing the requirement of LT activity of MltD for the suppression (Fig. 2D). Further, a *mltD* derivative lacking the C-terminal LysM domains severely reduced the suppression suggesting that the interaction of MltD with PG might be required for MltD activity (Fig. 2D). Overexpression of MltD was also able to modestly abrogate the vancomycin-sensitivity of *mepS* and *mepM* deletion mutants (Fig. 2E).

**Figure 2.**
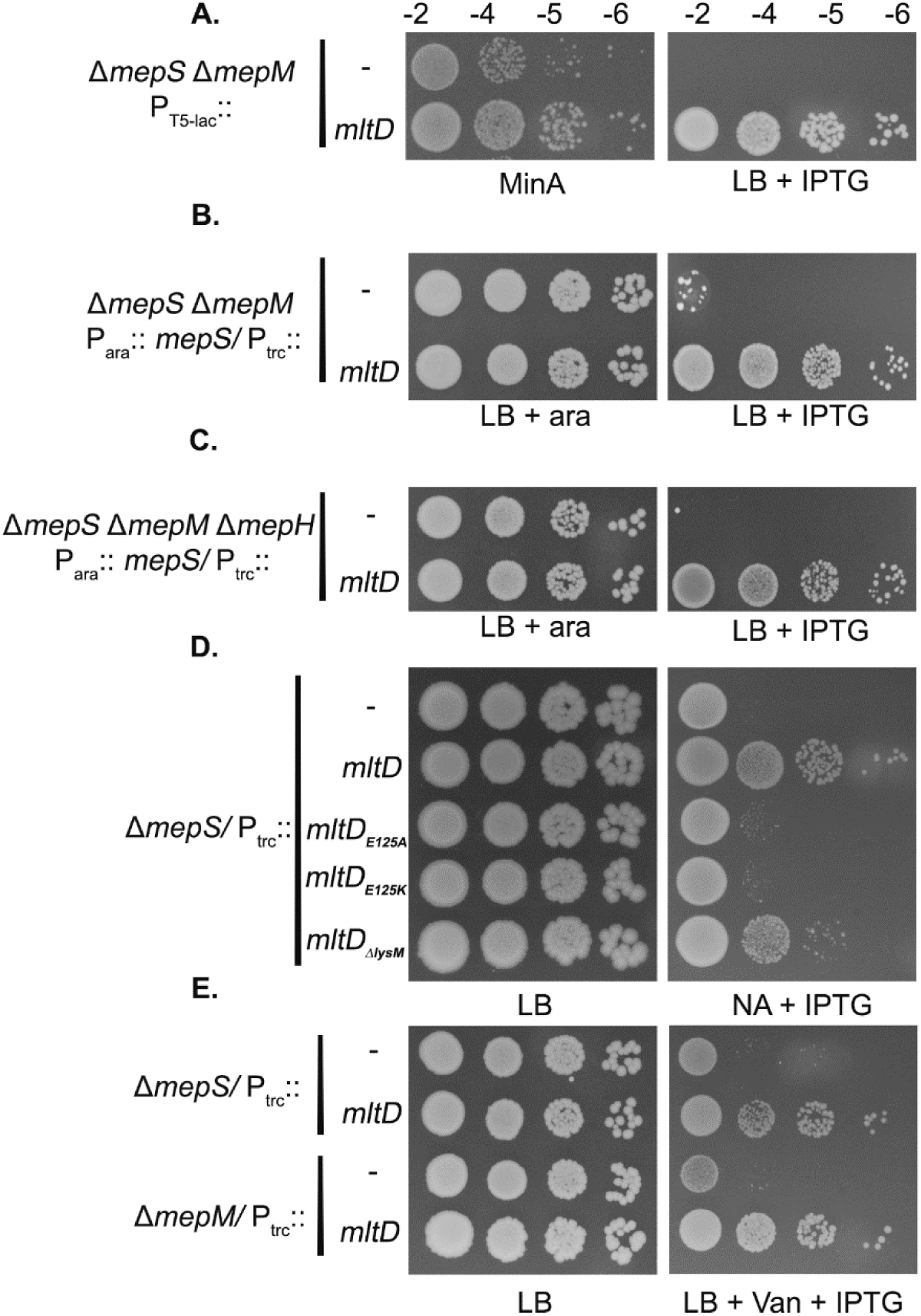
MltD overexpression compensates the deficiency of elongation-specific PG endopeptidases. (A) Cells of *mepS*::*frt mepM*::Kan carrying pCA24N (ASKA empty vector) or pCA24N-*mltD* were grown overnight in Min-A medium and viability was tested on LB plates with IPTG (25 µM). (B) Indicated mutant strains carrying pTrc99a vector or pTrc99-*mltD* were grown in LB with 0.2% arabinose and viability was checked on indicated plates. IPTG was used at 75 µM. (C) Growth of indicated strains with pTrc99a or pTrc-*mltD* was tested on LB plates without and with IPTG (500 µM). (D) Strains carrying pTrc99 or its *mltD* derivatives were grown overnight and viability was tested on LB and NA plates containing IPTG (10 μM). (E) Growth of indicated strains was checked on LB plates supplemented with and without IPTG (25 μM) having 225 µg/ml vancomycin.

Further, to modulate the expression of MltD, we constructed a strain in which the native *mltD* promoter is replaced by a *lac* promoter at its chromosomal locus (as described in SI). As expected, this construct rescued the growth defects of *mepS* deletion mutant in an IPTG-dependent manner (Fig. S1E). All these observations collectively indicated that overexpression of MltD, a LT, is able to suppress the growth defects caused by absence of the essential cross-link cleaving endopeptidases, MepS, MepM and MepH.

### MltD works in conjunction with MepS

Because MltD overexpression rescued the growth of the elongation-specific endopeptidase deletion mutants, we examined the role of MltD in PG expansion. MltD deletion mutant by itself did not exhibit any discernible growth defect on LB, LBON or NA at any temperature or sensitivity to cell-wall antibiotics such as cefsulodin, mecillinam or vancomycin. However, deletion of *mltD* exacerbated the NA-sensitive phenotype of *mepS* mutant (Fig. 3A, 3B) and also hampered the growth of *mepS mepM* double mutant on minimal media (Fig. 3C). To further examine the role of MltD in PG expansion, we measured the level of nascent PG strand incorporation using tritiated mDAP [27] in *mltD*, *mepS*, *mepS mltD* and *mepM mltD* mutants. Fig. 3D shows that absence of MltD further reduces the rate of PG incorporation in *mepS* mutant but had no significant effect either in the WT or in the *mepM* mutant. The above results suggest that MltD collaborates with MepS for optimal PG synthesis raising an interesting possibility of glycan strand cleavage contributing to the expansion of PG.

**Figure 3.**
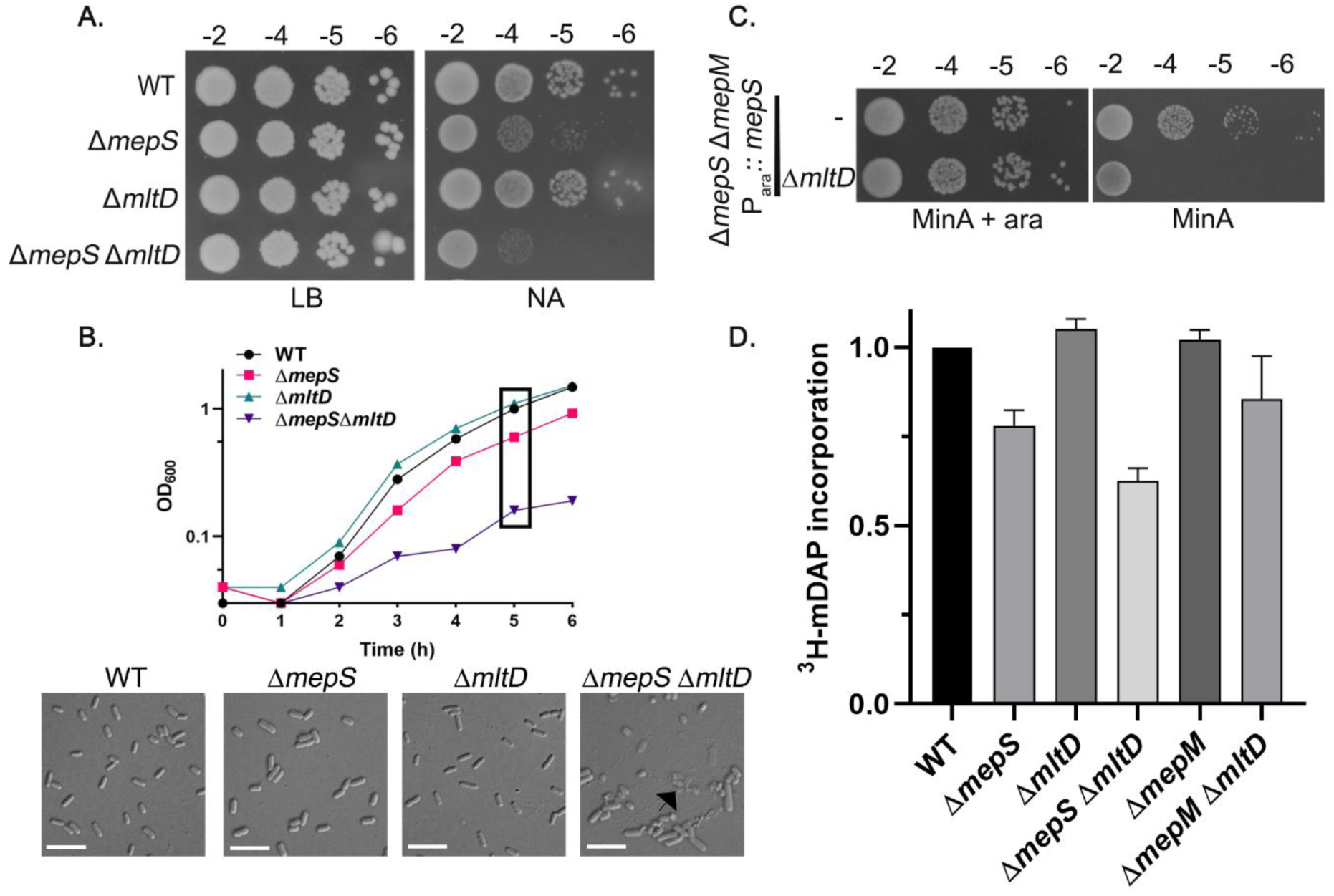
Absence of MltD exacerbates the MepS^−^ phenotypes. (A) WT and its mutant derivatives were grown overnight in LB and 4 µl was spotted on LB and NA plates and grown at 30°C. (B) Overnight grown cultures of the aforementioned strains were washed and diluted 1:100 into fresh NB at 30°C and growth was monitored by OD_600_. Cells were collected after 5 h of growth (as indicated) and subjected to DIC microscopy as described in Materials and Methods. Arrow indicates lysed cells. Scale bar is 5 µm. (C) Indicated strains or its derivatives with a deletion of *mltD* were grown with 0.2% arabinose overnight and the viability was checked on Min-A with and without arabinose supplementation (0.2%). (D) Bar diagram depicting the incorporation of ^3^H-mDAP into the PG sacculi of WT and its mutant derivatives. All strains had *lysA*::Kan deletion to block the conversion of mDAP into lysine. Strains grown overnight in LB were washed and sub-cultured in Min-A medium supplemented with 0.5% CAA and grown till OD_600_ of 0.4 to 0.5. Normalised cell fractions were immediately pulsed with 5 µCi/ml of ^3^H-mDAP for 10 min and processed as described in Materials and Methods [27]. Value of 1 corresponds to approximately 12,000-15,000 CPM. It is to be noted that *mltD* deletion mutant showed a consistent and a small increase of 5% in ^3^H-mDAP incorporation compared to that of WT.

### MltD is a substrate of NlpI-Prc proteolytic system

The results obtained thus far indicated that MltD is most likely regulated by NlpI-Prc system, prompting us to examine its level in strains lacking the components of NlpI-Prc. For this, we constructed a functional C-terminal 3X Flag-tagged MltD at its native chromosomal locus (as described in SI) and examined its levels in WT, *nlpI*, *prc,* and *nlpI prc* mutants. Fig. 4A shows that MltD level is approximately 2.5-fold higher in *nlpI* and *prc* single mutants or in *nlpI prc* double mutant compared to that of WT indicating that both NlpI and Prc together contribute to the degradation of MltD. In addition, MltD exhibited growth-phase specificity with its levels high till the late-exponential phase of growth (up to an OD_600_ of 2.0) and then declining to almost half as cells enter the stationary-phase (Fig. 4B). Interestingly, MltD exhibited similar pattern of growth-phase specificity even in the absence of NlpI-Prc system indicating another layer of regulation that may operate to control the MltD level in stationary-phase (Fig. 4B). In addition, spectinomycin-chase experiments showed that half-life of MltD in absence of NlpI-Prc is comparable to that of the WT confirming the existence of an alternate proteolytic system that degrades MltD (Fig. S2A). To examine whether other OM-associated LTs are also regulated similarly by NlpI-Prc system, we checked the cellular levels of two other LTs, MltA (*mltA*-Flag) and MltF (*mltF*-Flag) in the absence of NlpI or Prc. Interestingly, MltF levels were approximately two-fold higher in absence of Prc but not in the absence of NlpI whereas MltA level is not affected by either of them (Fig. S2B). It is known earlier that the levels of three other LTs-MltB, MltG and DigH are modulated by the Prc protease [21,28].

**Figure 4.**
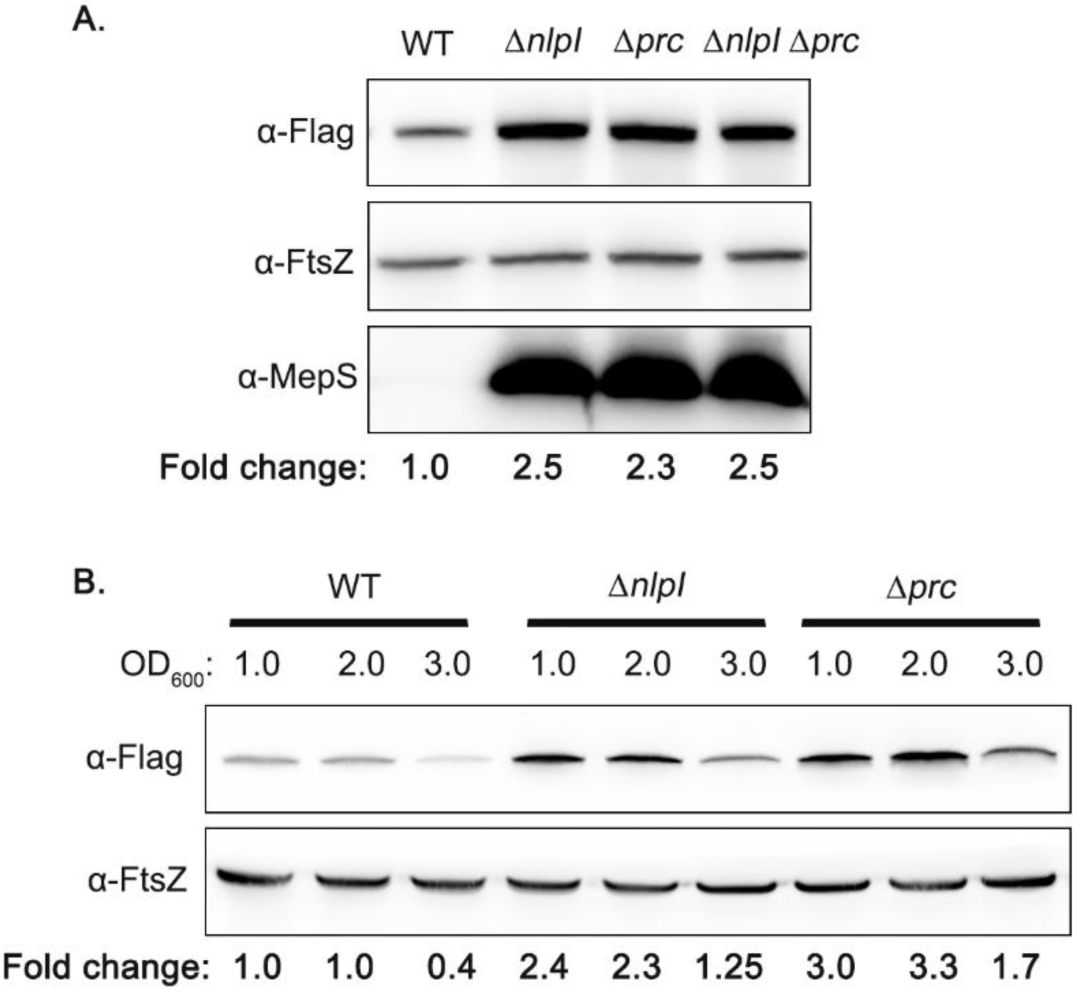
MltD is a substrate of NlpI-Prc proteolytic system. (A) Indicated strains carrying *mltD*-Flag at their native chromosomal locus were grown in LB and fractions were collected between OD_600_ of 0.8 -1.0. The fractions are normalized to 1.0 OD, cell lysates were made, and aliquots equivalent of 0.5 OD cells were subjected to SDS-PAGE followed by western blotting. MepS is used as a positive control. FtsZ is used as loading control to normalize the target protein. Bands were quantitated using ImageJ software. Fold-change was calculated after normalization with the FtsZ values. All the experiments were done three times and the representative blots are shown. (B) Western blot showing the growth-phase specific expression of MltD-Flag in the indicated strains. Cells were grown in LB and fractions were collected at different OD_600_ (1.0, 2.0 and 3.0) values and processed as described above.

### MltD is regulated by stationary-phase specific sigma factor, RpoS

Because the above observations suggested stationary-phase specific control of MltD, we examined the effect of σ^S^ (RpoS), an alternative stationary-phase specific sigma factor [29] on MltD stability. Fig. 5A shows ∼3-fold higher MltD in the absence of RpoS exclusively in the stationary-phase cells of *E. coli*. RpoS is a subunit of RNA polymerase that influences transcription of several genes during growth of *E. coli* in various stress conditions including starvation, stationary-phase, and osmolarity [29, 30]. To check the effect of RpoS on *mltD* gene expression, we constructed a chromosomal *lacZ* fusion downstream to the promoter of *mltD* (P_mltD_::*lacZY*; as described in SI) [31] and measured β-galactosidase levels. However, the *lacZ* levels remained identical in both WT and *rpoS* deletion strain suggesting that RpoS does not regulate *mltD* at the step of transcription (Fig. S3A). To examine this further, we constructed a plasmid carrying an *mltD*-Flag gene (lacking its native promoter) downstream to an IPTG-inducible promoter (P_trc_::*mltD*-Flag) and checked MltD levels in the WT and *rpoS* mutant. Fig. 5B shows that the plasmid-borne MltD level is highly elevated in the absence of RpoS indicating its control on MltD is promoter-independent and post-transcriptional. To check the possibility of RpoS modulating MltD at the level of post-translational stability through upregulation of other periplasmic proteases, we deleted genes encoding crucial periplasmic proteases (DegP, DegQ and PtrA) and examined MltD levels. However, MltD levels in the absence of these proteases remained the same as that of the WT indicating RpoS does not govern MltD levels through these proteases (Fig. S3B). Overall, these observations suggest RpoS exerts its control on MltD either at the step of translation or post-translational stability.

**Figure 5.**
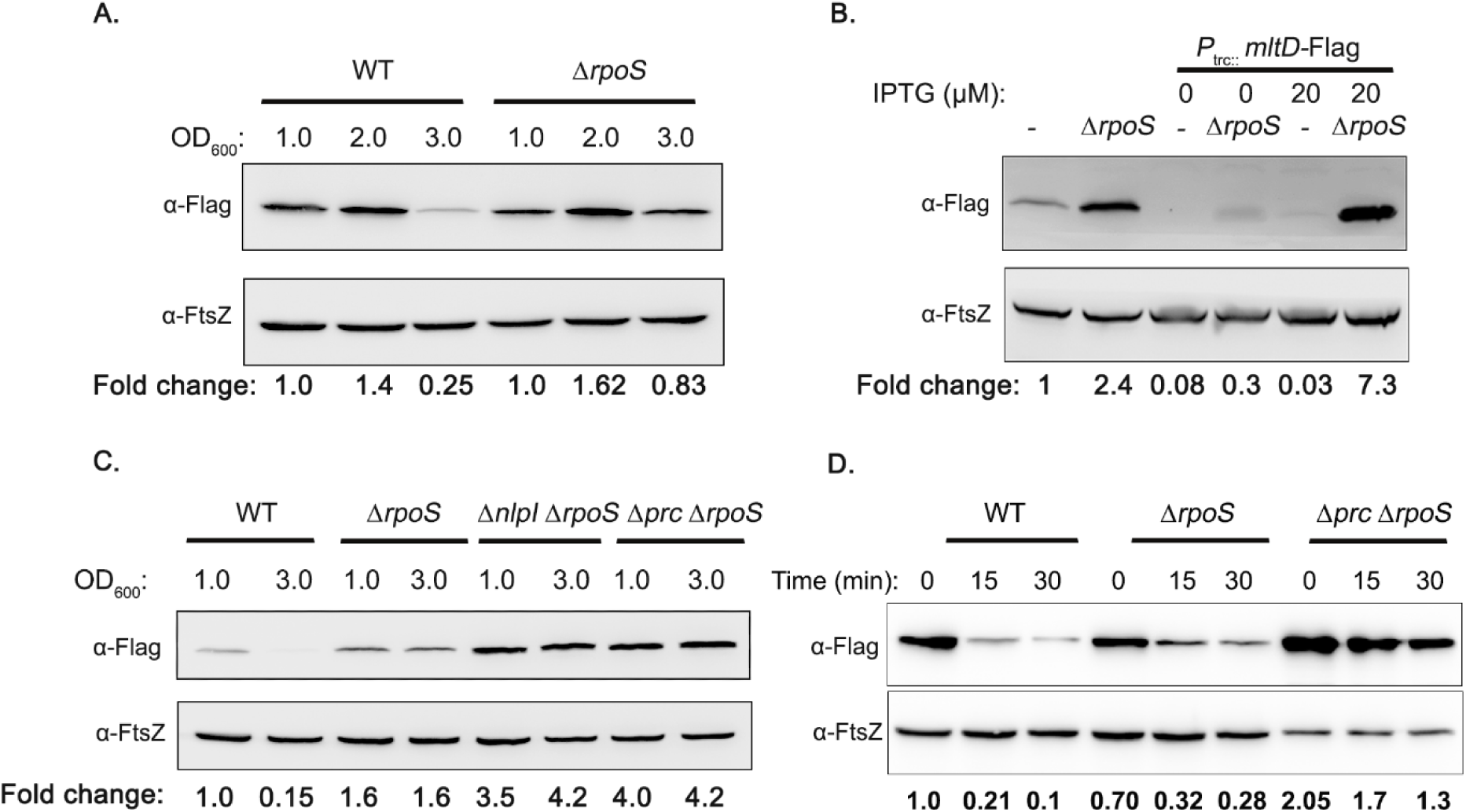
Regulation of MltD by stationary phase specific sigma factor, RpoS. (A) Indicated strains were grown in LB and fractions were collected at different OD_600_, and MltD levels were analysed by western blotting (B) WT and Δ*rpoS* mutant strain carrying either chromosomal MltD-Flag or a plasmid-borne MltD-Flag (P_trc::_*mltD*-Flag) were grown in LB till OD_600_ of 3 and cell lysates equivalent of 0.5 OD were subjected to SDS-PAGE followed by western blotting. IPTG was used at 20 μM. (C) Indicated strains were grown in LB and cell fractions were collected at OD_600_ of 1.0 and 3.0. Normalised cell fractions were used for western blot analysis. (D) Half-life of MltD-Flag in the indicated strains was checked as follows: cells were grown in LB and 300 µg/ml of spectinomycin was added to block translation at an OD_600_ of 1.0. Fractions were collected at indicated time points and analysed by western blotting.

Next to check whether NlpI-Prc system and RpoS act independently, we constructed double deletion mutants, *nlpI rpoS* and *prc rpoS* and checked the levels of MltD in both exponential and stationary phase (Fig. 5C). We find that in these strain backgrounds, the MltD levels remain highly stabilized all through the growth cycle indicating that NlpI-Prc and RpoS regulatory systems work in parallel to stringently control the cellular levels of MltD. The spectinomycin chase experiments show that MltD is highly stabilized in absence of both these systems (Fig. 5D) indicating these two pathways play a major role in the maintenance of cellular MltD levels.

### MltD is degraded by NlpI-Prc in vitro

So far, our results indicated that MltD is a substrate of NlpI-Prc proteolytic machinery *in vivo*. To test this observation *in vitro*, we overexpressed and purified MltD, NlpI and Prc lacking their signal sequences as hexahistidine fusion proteins. We then performed *in vitro* degradation assay by co-incubating these proteins and checked by SDS-PAGE (Fig. 6; Fig. S4A). Fig. 6 shows that MltD degradation by the protease Prc is very rapid in the presence of NlpI (lanes 4-8) whereas co-incubation of MltD with Prc protease in absence of NlpI does not lead to its degradation (lanes 2-3).

**Figure 6.**
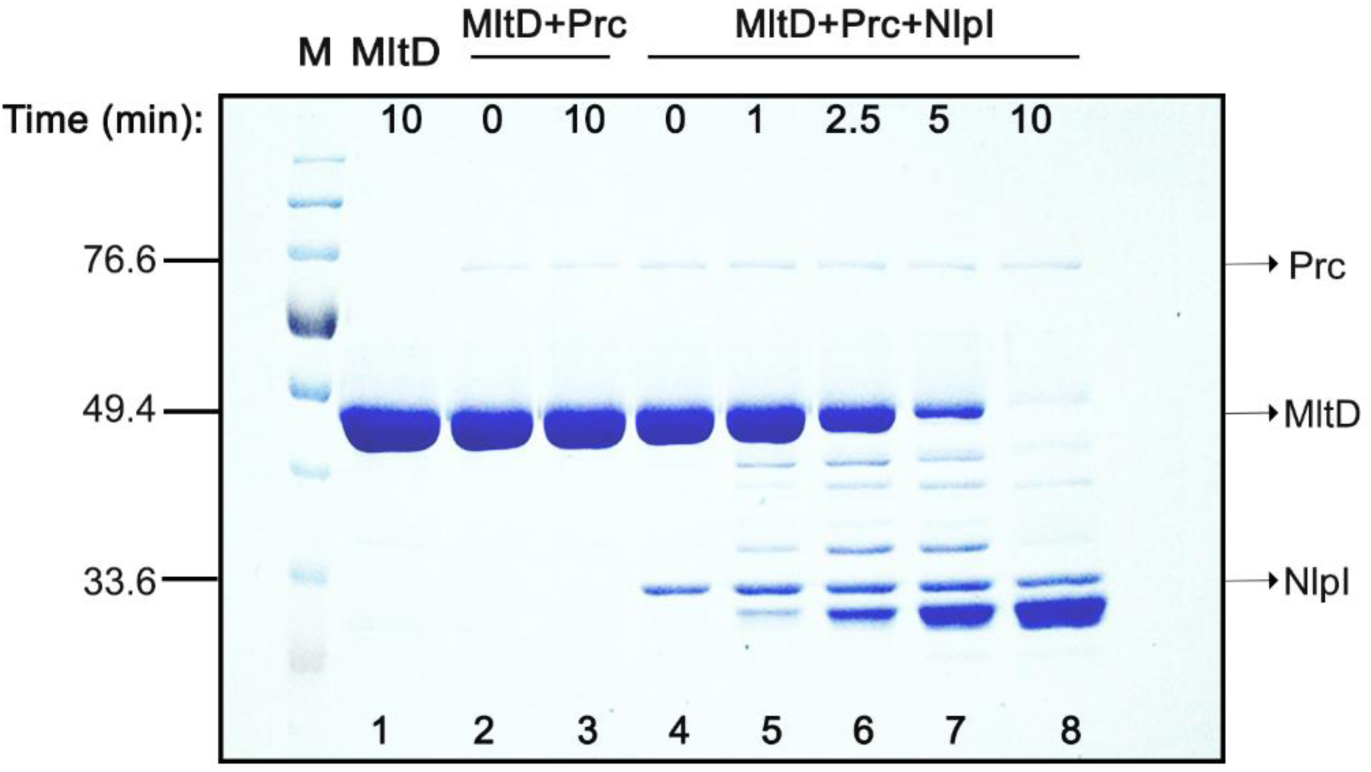
MltD degradation by Prc is facilitated by NlpI. *In vitro* degradation assay with purified MltD, Prc and NlpI proteins. The proteins were mixed in different combinations and incubated at 37°C as indicated. They were subjected to SDS-PAGE followed by Coomassie brilliant blue staining. The following amount of proteins were used: MltD-10 µg, NlpI-1 µg and Prc-0.4 µg. Molecular mass of the proteins is indicated in kDa.

Both *in vitro* and *in vivo* experiments showed that MltD is degraded by the Prc protease, and this degradation is contingent on the presence of its adaptor protein, NlpI indicating the turnover of MltD is regulated by NlpI-Prc system. Previous studies have shown that the turnover of MepS is similarly controlled by the NlpI-Prc [12]. Therefore, to check whether MltD or MepS is the preferred substrate of NlpI-Prc proteolysis, we purified MepS (as described in SI) and performed degradation assays either individually or in combination with MltD (Fig. S4B). However, preliminary experiments indicate both MltD and MepS are degraded by the NlpI-Prc system to a comparable extent (Fig. S4B).

### MltD activity is inhibited by NlpI

Considering an earlier study [32] which showed NlpI binding and modulating the activity of several PG endopeptidases, we examined whether NlpI can similarly bind and/ or alter the activity of MltD. For this purpose, we prepared PG sacculi from WT *E. coli* cells (MG1655) and treated the sacculi with purified MltD. After digestion, the soluble muropeptides were separated by RP-HPLC (reverse phase-high pressure liquid chromatography) and the chromatograms were analysed (Fig. 7C). Next, to check whether addition of NlpI affects the enzymatic activity of MltD, an equimolar amount of NlpI was pre-incubated with MltD for 10 min and subsequently, this protein mixture was used to treat the PG sacculi (Fig. 7D). The chromatograms show that addition of NlpI significantly decreases the enzymatic activity of MltD. More importantly, the reduction in the activity of MltD by NlpI was specific to MltD (Fig. 7C, D); the activity of a soluble periplasmic LT, Slt70, remained unchanged in the presence of NlpI (Fig. 7A, B).

**Figure 7.**
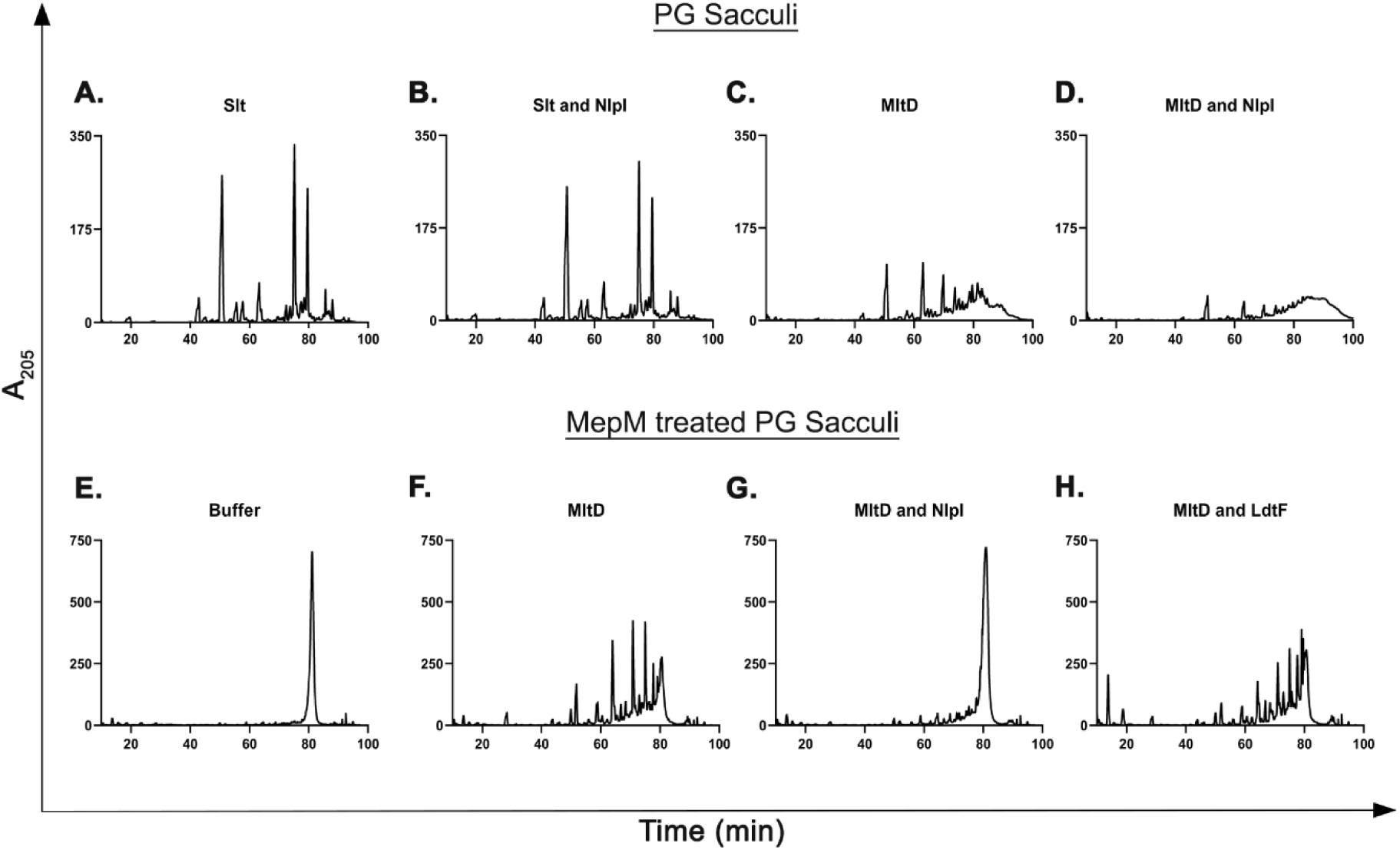
Enzymatic activity of MltD and its inhibition by NlpI. HPLC chromatograms showing the activity of LTs (Slt and MltD) on PG sacculi. Sacculi were treated with either Slt or MltD (panel A,C) or treated with Slt+NlpI or MltD+NlpI (panel B,D) for 16 h at 37°C. Slt or MltD were preincubated with NlpI for 10 min prior to treatment. To generate un-crosslinked PG substrates, the sacculi were initially treated with MepM, a known 4-3 cross-link specific endopeptidase for 16 h at 37°C followed by another 16 h of treatment with indicated proteins (E,F,G,H). Addition of NlpI but not LdtF [33] inhibited the activity of MltD. All proteins were used at 5 µM.

Earlier studies indicated MltD may prefer un-crosslinked glycan strands as its substrate [18]. Therefore, we generated un-crosslinked PG substrates by treating the PG sacculi initially with a D,D-endopeptidase MepM which cleaves the 4,3 cross-links between the glycan strands [6]. The MepM-treated PG sacculi were then used for digestion with MltD and the chromatograms show that MltD is able to hydrolyse the uncross-linked PG with higher efficiency compared to that of intact PG sacculi (Fig. 7C vs 7F). Consistent with the above observations, addition of NlpI severely reduced the hydrolytic activity of MltD on uncross-linked PG (Fig. 7G). Addition of another PG endopeptidase, LdtF [33] did not have any effect on the activity of MltD and served as a negative control (Fig. 7H).

### Interaction of MltD with NlpI could be transient

We have thus far observed that MltD and MepS show similarity in terms of their growth-phase specificity and also the ability to be controlled by NlpI-Prc proteolytic system. In addition, MltD activity was hindered in the presence of NlpI. Therefore, we wondered whether MltD physically interacts with NlpI and/ or MepS to form a large multi-protein hydrolase complex to facilitate PG expansion. To test this idea, we performed a co-purification (*in vivo* pull-down assay) using a plasmid-borne NlpI-His as a bait to examine the NlpI-MltD interaction (Fig. S4C). However, we observed that NlpI-His did not pull down MltD, although it was able to pull down MepS and Prc as shown earlier [12]. We further checked whether MepS and MltD form a complex using plasmid-borne MepS-His as a bait but did not find any interaction between them either in the WT strain or in the absence of NlpI (where MltD is expected to be in higher amount) suggesting NlpI may transiently interact with MltD (Fig. S4D).

## DISCUSSION

In this study, we demonstrate that cleavage of glycan polymers by a lytic transglycosylase, MltD plays a role in cell wall expansion. It is known that cutting the PG mesh by hydrolysis of cross-links between the glycan strands is crucial for its expansion; and here, we identify a previously unknown role for the glycan strand cleavage in opening the mesh to generate space for the expansion of PG in *E. coli*. We also find that MltD is stringently controlled by two independent regulatory mechanisms. It is controlled at the step of post-translational stability by NlpI-Prc mediated proteolysis and in addition, the MltD abundance in the stationary-phase cells is governed by RpoS, a stationary-phase specific sigma factor, highlighting the need for the maintenance of optimal level of MltD during growth of *E. coli*.

### Role of MltD in PG expansion

MltD is one of the several membrane-bound LTs encoded by *E. coli* [10,18,34]. It is an OM lipoprotein containing an N-terminal transglycosylase domain and two LysM (lysin) repeats which facilitate PG binding (Fig. 1D) [25]. It is reported to be an endolytic LT that may prefer uncross-linked substrates; however, its role in PG metabolism is not clear [17]. Our initial finding of MltD stabilization as the basis of suppression of *mepSM* double mutant in absence of NlpI-Prc indicated a potential role for MltD in PG expansion (Fig. 1). Subsequently, overexpression of MltD was shown to suppress several growth phenotypes of *mepS*, *mepM*, *mepH* mutants (Fig. 2). Notably, overexpression of MltD compensating the absence of all the three essential cross-link specific endopeptidases indicated that opening the mesh by endolytic cleavage of the glycan polymers may also facilitate PG expansion.

The additive growth phenotypes of *mepS* and *mltD* mutants (Fig. 3) suggest that under normal physiological conditions these two hydrolases collaborate for optimal PG expansion. Here, we speculate that MepS may initially cleave the peptide cross-links followed by cleavage of the uncross-linked glycan strands by MltD leading to generation of a gap in the PG for incorporation of new strands (Fig. 8). Interestingly, the turnover of both MepS and MltD is controlled by the NlpI-Prc system, suggesting the existence of a shared regulatory pathway that responds to the cellular need of PG expansion. During cell elongation, their stabilization leads to PG hydrolysis to accommodate nascent PG strands whereas, when PG synthesis is not needed, MepS and MltD may undergo NlpI-Prc dependent proteolysis thus avoiding a futile cycle of PG degradation (Fig. 8). The gap thus generated by the hydrolases may allow interaction of the IM-localized aPBPs with their cognate Lpo activators located in the OM subsequently activating the synthesis and formation of cross-links to expand the PG sacculus [3,4].

**Figure 8.**
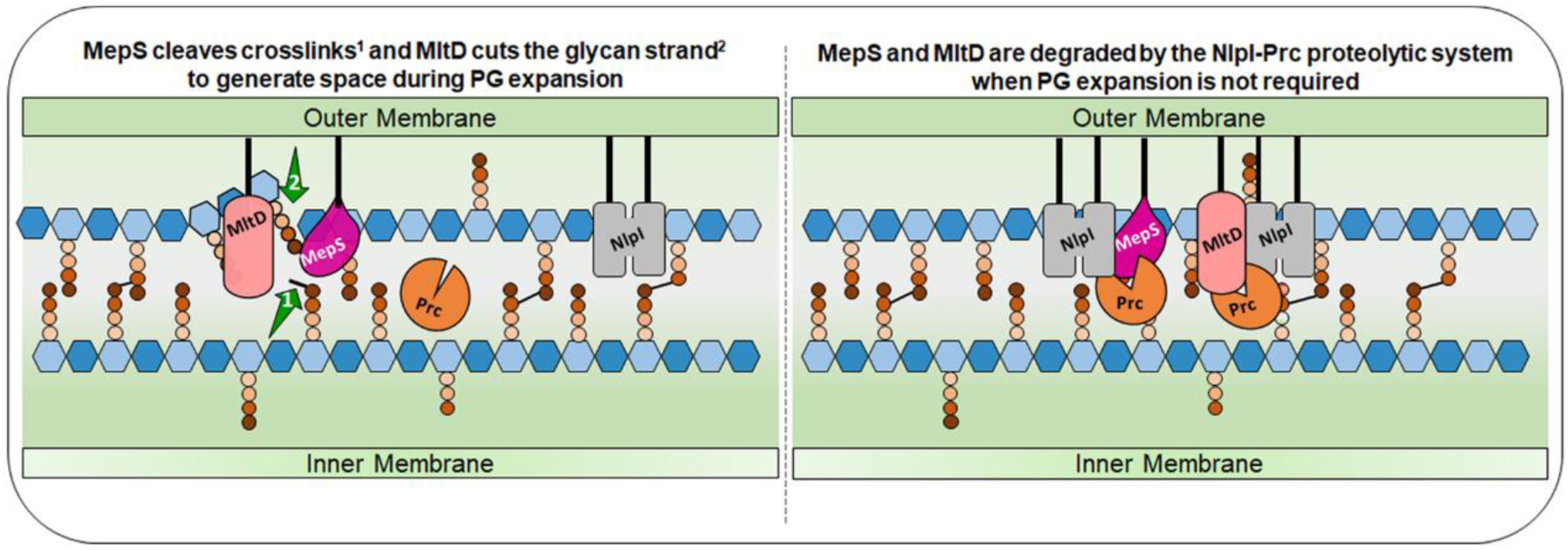
Model depicting the role of MltD and MepS in PG expansion. During PG expansion, activity of a key endopeptidase, MepS and the lytic transglycosylase activity of MltD generate space for the incoming PG material (left panel). Both these hydrolases may work in tandem for efficient cleavage of the PG sacculi. Since MltD prefers uncross-linked PG, we speculate MepS could initiate the cross-link hydrolysis (indicated by a green arrow^1^) followed by glycan strand cleavage by MltD (indicated by green arrow^2^). When PG expansion is not required, both MepS and MltD are degraded by the NlpI-Prc proteolytic system thereby avoiding a futile cycle of PG degradation (right panel).

### Regulation of MltD

Although PG hydrolases are fundamental for various cell wall processes, they need to be tightly regulated as their unfettered activity leads to catastrophic PG degradation with loss of envelope integrity. In support of this, we find MltD levels are controlled by two independent pathways. Both *in vivo* and *in vitro* experiments confirmed MltD to be a substrate of NlpI-Prc proteolytic system (Fig. 4; Fig. 6). Till date, MepS [12] and MepH [11], the cross-link specific endopeptidases involved in PG enlargement are the only substrates identified to be dependent on NlpI for their turnover by Prc. In contrast, Prc protease alters the stability of several periplasmic PG enzymes in an NlpI-independent manner [15,28]. The role of MepS, MepH and MltD in PG expansion, and their control by NlpI-Prc system raises an interesting possibility of NlpI playing a larger role in regulating the process of PG expansion by modulating the stability of the space-making PG hydrolases. Interestingly, unlike MepS (whose degradation is solely dependent on NlpI-Prc) [12], MltD is partly controlled by NlpI-Prc system (Fig. 4B, Fig. S2A) signifying MltD’s role in other PG-related processes. Although MltD single deletion mutant has no phenotype, absence of MltD along with other LTs leads to cell-chaining phenotype implying its role in daughter cell-separation [35].

In addition to NlpI-Prc control which may operate during cell wall expansion, MltD levels are kept in check in stationary-phase by RpoS, the stationary-phase specific sigma factor (Fig. 5); although our results show that the regulation is post-transcriptional, it is not yet clear how RpoS mediates MltD degradation. We speculate RpoS may increase the transcription of a periplasmic protease in stationary phase cells of *E. coli* which then partakes in the degradation of MltD.

### Role of LTs in PG metabolism of *E. coli*

*E. coli* encodes several redundant LTs reflecting their importance in various cellular processes [16,18,24]. However, a mutant lacking LTs has no major phenotype, hindering the progress in understanding their specific functions. Recent studies have now revealed their distinct roles in PG pathways [23,24,34]. For example, the soluble LT, Slt70 cleaves the uncross-linked glycan strands during PG synthesis [22]; MltG determines the glycan chain length during PG polymerization [20]; and DigH cleaves the denuded glycan strands at the cell septum [21]. To date, cross-link cleavage by endopeptidases is believed to facilitate sacculus expansion, however, our study shows a role for glycan strand cleavage, mediated by MltD in PG sacculus expansion.

## MATERIALS AND METHODS

### Media and growth conditions

Strains were normally grown in Luria-Bertani (LB) (1% tryptone, 0.5% yeast extract and 1% NaCl) broth. LBON is LB without NaCl. Nutrient Agar (NA) had 0.5% peptone, 0.3% beef extract and 1.5% agar. Minimal-A medium [36] was supplemented with B1, MgS0_4_ and 0.2% glucose before use. Casamino acids (CAA) were used at 0.5%. Antibiotics were used as following: kanamycin (Kan)-50 µg/ml; chloramphenicol (Cm)-25 µg/ml; ampicillin (Amp)-50 µg/ml. Isopropyl β-D-thiogalactopyranoside (IPTG) and L-arabinose were used at the indicated concentration. Growth temperature was 37°C unless otherwise specified. Growth was monitored by measuring optical density at 600nm (OD_600_).

### Molecular and genetic techniques

Recombinant DNA techniques, P1-phage mediated transductions and transformations were performed using standard methods [36]. Deletion mutations were sourced from Keio mutant collection [37]. Whenever required, the kanamycin resistance gene was flipped out using pCP20 plasmid encoding FLP recombinase [38]. The 3X Flag epitope tagging was done at the C-terminus of the ORF at its native chromosomal locus by recombineering [39].

### Viability assays and microscopy

Viability of strains was tested by serially diluting (10^-2^, 10^-4^, 10^-5^ and 10^-6^) the overnight-grown cultures and applying 4 µl of each dilution onto the indicated plates followed by incubation for 18-36 h at the indicated temperature. Growth was recorded by photographing the plates. For microscopic analysis, overnight cultures were subcultured and grown till the required OD. Cells were collected, immobilized onto a thin 1% agarose pad and visualized using a Zeiss apotome microscope by DIC (Nomarski optics).

### Plasmid constructions

For PCR amplifications, genomic DNA of MG1655 strain was used as a template. Amplification of DNA was done using Q5 High-Fidelity DNA polymerase (NEB) and clones obtained were confirmed by sequence analysis.

Plasmid constructions are described in detail in SI.

### Screen for isolation of spontaneous suppressors of *mepS mepM* double mutant

A strain carrying deletion of both *mepS* and *mepM* was grown overnight in minimal-A broth and the next day, approximately 10^8^ cells were plated on LB agar and incubated at 37°C. Approximately 30 colonies that were able to grow on LB plates were picked up and purified. The suppressors were broadly classified based on the extent of suppression and a few of these were initially mapped using conventional conjugational and transductional mapping techniques [36]. Interestingly, these suppressor mutations turned out to be recessive alleles of either *nlpI* or *prc*; subsequently we tested all the remaining suppressors and to our surprise all the short-listed suppressors mapped to either *nlpI* or *prc*.

### Western blotting

For collection of samples for the western blot analysis, overnight cultures were subcultured and grown till indicated OD_600_ value. Growth was carefully monitored by measuring the culture density (OD_600_) at regular intervals and cells were collected. Normalized cell fractions were immediately pelleted at 4⁰C, resuspended in 1X Laemmli loading buffer and boiled for 10 min. Samples were centrifuged before being resolved using SDS-PAGE (14%). Proteins were transferred onto a PVDF membrane using semi-dry western transfer method. Bands were visualized by Ponceau S stain to ascertain protein transfer. Membrane was blocked by 5% skimmed milk in 1X TBST for 1 h and then incubated overnight with primary antibodies [1:3,000 for α-MepS (kind gift from Waldemar Vollmer), 1:5,000 for α-NlpI (laboratory collection), 1:4,000 for α-Flag (F3165, Sigma-Aldrich, USA), 1:10,000 for α-FtsZ (Kind gift from Joe Lutkenhaus), at 4°C under constant shaking. The membrane was then washed four times with 1 X TBST (Tris-buffered saline with 0.01% Tween-20) for 5 min each and then probed with secondary antibodies (dilution of 1:10,000) tagged to HRP (horseradish peroxidase) enzyme and further incubated for 1 h. Membrane was then washed with 1X TBST four times for 5 min each to remove unbound antibodies. Proteins were visualized using enhanced chemiluminescence (ECL)-prime detection substrate (Amersham) inside a chemi-documentation system. Bands were quantitated using ImageJ software. FtsZ is used as loading control to normalize the target protein. Fold-change was calculated after normalization with the FtsZ values. All the experiments were done three times and the representative blots are shown.

### Half-life determination

To examine the MltD protein degradation *in vivo*, spectinomycin-chase experiments were performed. To rapidly growing cells in LB (at the required OD), spectinomycin (300 µg/ml) was added to inhibit the protein synthesis and cells were collected at indicated time points. Normalized cell fractions were processed by western blotting as described above.

### mDAP incorporation assay

Incorporation of ^3^H-mDAP (tritiated mDAP) into the PG sacculi was done as described [27]. Indicated strains (lacking LysA to block the formation of lysine from mDAP) were grown overnight in LB broth, and the next morning, they were washed, and diluted 1:100 in prewarmed minimal-A medium containing 0.2% glucose and 0.5% CAA. At an OD of 0.3-0.4, normalized cell fractions were collected and incubated with 5 µCi/ml of ^3^H-mDAP (^3^H-DAP;14.6 Ci mmol-1, Moravek Biochemicals, USA) for 10 min with gentle shaking at 37°C. Cells were immediately lysed by addition of 3 ml of 4% SDS and boiled for 1 h. The mixture was cooled overnight at RT and filtered through 0.22 µm filter (Millipore). The insoluble PG sacculi were collected on the filters, washed with 30 ml of Milli-Q water and the filters were dried completely before counting the radioactivity in a liquid scintillation counter (Perkin-Elmer).

### Protein overexpression and purification

MepS, MltD, NlpI, Prc, Slt, and MepM proteins (lacking their periplasmic signal sequences) fused to hexahistidine tags were overexpressed and purified to homogeneity from BL21 (λDE3) or its derivatives as described in detail in SI. MltD and MepS had N-terminal His tag whereas NlpI, Prc, Slt and MepM were fused to C-terminal His tag. Proteins were purified through metal-affinity chromatography using Ni^2+^–NTA agarose beads (Qiagen). Purified protein aliquots were stored at -30°C in 50 mM Tris-Cl, 100 mM NaCl, 1mM DTT and 50% glycerol for further use.

### *In vitro* degradation assay

The indicated purified proteins were mixed and incubated at 37°C for different time intervals as described earlier [12]. Reactions were stopped by addition of 4X Laemmli buffer and boiling for 10 min. Samples were separated by SDS-PAGE followed by staining with Coomassie brilliant blue. Each reaction contained 10 µg MltD, 1 µg NlpI and 0.4 µg Prc. Proteins were added and mixed in the reaction buffer (20 mM Tris-Cl pH 8.0, 150 mM NaCl) on ice.

### Peptidoglycan isolation

PG sacculi were isolated as described earlier with slight modifications [2,6]. In brief, cells from 1 L of exponentially growing culture of WT (MG1655) were harvested by centrifugation followed by resuspension of the cell pellet in 9 ml of ice-cold deionised water. The cell suspension was added drop wise to 9 ml of boiling 8% SDS with continuous stirring and further boiled for 60 min. This mixture was stored overnight at RT and the sacculi were collected by ultracentrifugation (400,000 g for 60 min at RT). The pellet was washed repeatedly with deionised water to remove SDS. The high molecular weight glycogen and covalently bound lipoprotein were removed by adding α-amylase (100 µg/ml in 10 mM Tris-HCl, pH 7.2 for 2 h at 37°C) and heat-activated pronase (200 µg/ml for 90 min at 60°C) respectively. Samples were boiled with equal volume of 8% SDS for 15 min to inactivate these enzymes. Sacculi were collected by ultracentrifugation and washed repeatedly with deionized water to remove traces of SDS. Finally, the pellet was resuspended in 25 mM Tris-HCl (pH 8.0) and stored at -30°C.

### PG analysis by RP-HPLC

HPLC analysis was performed as described earlier [2,6] with some modifications. PG sacculi were treated with appropriate enzymes for 16h at 37°C, the proteins were heat-inactivated through boiling, and insoluble material was removed by centrifugation (14,000 RPM at RT). Subsequently, the soluble muropeptides were collected and mixed with equal volume of 50 mM sodium borate buffer (pH 9.0). The anomeric carbons of muropeptides were reduced by adding 1 mg of sodium borohydride. Excess borohydride was destroyed by adding 1/20^th^ volume of orthophosphoric acid and pH was adjusted between 2-4 before loading onto a Zorbax 300 SB RP-C18 (250 X 4.6mm, 5 µm) column. Separation of muropeptides was done by RP-HPLC using Agilent technologies RRLC 1200 system. Samples were injected onto a preheated column at 55°C and binding was allowed at a flow rate of 0.5 ml per minute with a solvent containing 1% acetonitrile and 0.1% trifluoroacetic acid (TFA) for 10 min. A gradient of 1-10% (MepM treated substrates) or 1-15% (untreated PG) acetonitrile containing 0.1% TFA was used for final elution at a flow rate of 0.5 ml per minute. Absorbance was detected at 205 nm.

### Supporting Information (SI)

**S1 Text** Details of strain and plasmid constructions, supplemental methods, and protocols are described.

**Figure S1:** Absence of NlpI-Prc system rescues the growth defects of *mepS mepM* mutant through MltD.

**Figure S2:** Regulation of LTs by NlpI-Prc proteolytic system.

**Figure S3:** Post-transcriptional control of RpoS on MltD.

**Figure S4:** Interaction between MltD, NlpI and Prc proteins.

**Table S1:** List of strains used in this study.

**Table S2:** List of plasmids used in this study.

## ACKNOWLEDGEMENTS

We thank NBRP (Japan): *E. coli* for Keio mutant collection and ASKA plasmid library; L SaiSree for construction of mutants used for genetic mapping; Krishna Kumari for help with HPLC, and members of Reddy lab for comments on the manuscript. This work is supported by funds from Council of Scientific and Industrial Research (MLP0141) and Department of Biotechnology (BT/PR33064/BRB/10/1819/2019) Government of India. SKM acknowledges financial support from CSIR.

## Author Contributions

MK and MR conceived the study, designed experiments, analyzed data and wrote the paper. MK and SKM performed the experiments and analyzed the data.

## Notes

### Competing Interest Statement

The authors have declared no competing interest.

## REFERENCES

1. Höltje J-V (1998) Growth of the stress-bearing and shape-maintaining murein sacculus of *Escherichia coli*. Microbiol Mol Biol Rev 62: 181–203.

2. Glauner B, Holtje JV, Schwarz U (1988) The composition of the murein of *Escherichia coli*. J Biol Chem 263: 10088–10095.

3. Egan AJF, Errington J, Vollmer W (2020) Regulation of peptidoglycan synthesis and remodelling. Nat Rev Microbiol 18: 446–460.

4. Garde S, Chodisetti PK, Reddy M (2021) Peptidoglycan: structure, synthesis, and regulation. EcoSal Plus 9: 629–646.

5. Tomasz A (1984) Building and breaking of bonds in the cell wall of bacteria-the role for autolysins. Microbial cell wall synthesis and autolysis. Elsevier pp.3–12.

6. Singh SK, Saisree L, Amrutha RN, Reddy M (2012) Three redundant murein endopeptidases catalyse an essential cleavage step in peptidoglycan synthesis of *Escherichia coli* K12. Mol Microbiol 86:1036– 1051.

7. Dörr T, Cava F, Lam H, Davis BM, Waldor MK (2013) Substrate specificity of an elongation-specific peptidoglycan endopeptidase and its implications for cell wall architecture and growth of *Vibrio cholerae*. Mol Microbiol 89:949–962.

8. Srivastava D, Seo J, Rimal B, Kim SJ, Zhen S, Darwin AJ (2018) A proteolytic complex targets multiple cell wall hydrolases in Pseudomonas. mBio 9e00972–18.

9. Vollmer W, Joris B, Charlier P, Foster S (2008) Bacterial peptidoglycan (murein) hydrolases. FEMS Microbiol Reviews 32:259–286.

10. Heijenoort JV (2011) Peptidoglycan hydrolases of *Escherichia coli*. Microbiol Mol Biol Rev. 75: 636–663.

11. Jeon WJ, Cho H (2022) A cell wall hydrolase MepH is negatively regulated by proteolysis involving Prc and NlpI in *Escherichia coli*. Front Microbiol 13: 878049.

12. Singh SK, Parveen S, SaiSree L, Reddy M (2015) Regulated proteolysis of a cross-link-specific peptidoglycan hydrolase contributes to bacterial morphogenesis. Proc Natl Acad Sci USA 112:10956–10961.

13. Su MY, Som N, Wu CY, Su SC, Kuo YT, Ke LC et al (2017) Structural basis of adaptor-mediated protein degradation by the tail-specific PDZ-protease Prc. Nat Comm 8: 1–13

14. Chakraborty D, Darwin AJ (2021). Direct and indirect interactions promote complexes of the lipoprotein LbcA, the CtpA protease and its substrates, and other cell wall proteins in *Pseudomonas aeruginosa*. J Bacteriol 203: 003932.

15. Sommerfield AG, Darwin AJ (2022) Bacterial carboxyl-terminal processing proteases play critical roles in the cell envelope and beyond. J Bacteriol 204: e00628–21.

16. Dik DA, Marous DR, Fisher JF, Mobashery S (2017) Lytic transglycosylases: concinnity in concision of the bacterial cell wall. Crit Rev Biochem and Molecular Biol 52: 503–542.

17. Lee M, Hesek D, Llarrull LI, Lastochkin E, Pi H, Boggess B, Mobashery S (2013) Reactions of all *Escherichia coli* lytic transglycosylases with bacterial cell wall. J Am Chem Soc 135: 3311–3314.

18. Scheurwater, E, Reid CW, Clarke AJ (2008) Lytic transglycosylases: Bacterial space-making autolysins. Int J Biochem Cell Biol 40: 586–591.

19. Park J.T, Uehara T (2008) How bacteria consume their own exoskeletons (turnover and recycling of cell wall peptidoglycan). Microbiol Mol Biol Rev 72: 211–227.

20. Yunck R, Cho H, Bernhardt TG (2016) Identification of MltG as a potential terminase for peptidoglycan polymerization in bacteria. Mol Microbiol 99: 700–718.

21. Yakhnina AA, Bernhardt TG (2020) The Tol-Pal system is required for peptidoglycan-cleaving enzymes to complete bacterial cell division. Proc Natl Acad Sci USA 117: 6777–6783.

22. Cho H, Uehara T, Bernhardt TG (2014) Beta-lactam antibiotics induce a lethal malfunctioning of the bacterial cell wall synthesis machinery. Cell 6:1300–1311.

23. Weaver AI, Alvarez L, Rosch KM, Ahmed A, Wang GS, van Nieuwenhze MS et al (2022) Lytic transglycosylases mitigate periplasmic crowding by degrading soluble cell wall turnover products. elife11:73178.

24. Weaver A, Taguchi A, Dorr T (2023) Masters of misdirection: peptidoglycan glycosidases in bacterial growth. J Bacteriol 205: 00428–22.

25. Bateman A, Bycroft M (2000) The structure of a LysM domain from *E. coli* membrane-bound lytic murein transglycosylase D (MltD). J Mol Biol 299: 1113–1119.

26. Kitagawa M, Ara T, Arifuzzaman M, Loka-Nakamichi T, Inamoto E, Toyonaga H et al (2005) Complete set of ORF clones of *Escherichia coli* ASKA library (a complete set of *E. coli* K-12 ORF archive): unique resources for biological research. DNA Res. 12(5): 291–299.

27. Wientjes FB, Pas E, Taschner PEM, Woldringh CL (1985) Kinetics of uptake and incorporation of meso-diaminopimelic acid in different *Escherichia coli* strains. J Bacteriol 164: 331–337.

28. Hsu PC, Chen CS, Wang S, Hashimoto M, Huang WC, Teng CH (2020) Identification of MltG as a Prc protease substrate whose dysregulation contributes to the conditional growth defect of Prc-deficient *Escherichia coli*. Front Microbiol 11:1–16.

29. Battesti A, Majdalani N, Gottesman S (2011) The RpoS-mediated general stress response in *Escherichia coli*. Ann Rev Microbiol 65: 189–213.

30. Weber H, Polen T, Heuveling J, Wendisch VF, Hengge R (2005) Genome-wide analysis of the general stress response network in *Escherichia coli*: σ^S^-dependent genes, promoters and sigma factor selectivity. J Bacteriol 187: 1591–1603.

31. Ellermeier CD, Janakiraman A, Slauch JM (2002) Construction of targeted single copy lac fusions using λ Red and FLP-mediated site-specific recombination in bacteria. Gene 290: 153–161.

32. Banzhaf M, Yau HC, Verheul J, Lodge A, Kritikos G, Mateus A et al (2020) Outer membrane lipoprotein NlpI scaffolds peptidoglycan hydrolases within multi-enzyme complexes in *Escherichia coli*. EMBO J 39:1–20.

33. Bahadur R, Chodisetti PK, Reddy M (2021) Cleavage of Braun’s lipoprotein, Lpp from the bacterial peptidoglycan by a paralog of L,D-transpeptidases, LdtF. Proc Natl Acad Sci USA 118: e2101989118.

34. Martinez-Bond EA, Soriano BM, Williams AH (2022) The mechanistic landscape of lytic transglycosylase as targets for antibacterial therapy. Curr Opin Struct Biol 77: 102480.

35. Heidrich C, Ursinus A, Berger J, Schwarz H, Höltje JV (2002) Effects of multiple deletions of murein hydrolases on viability, septum cleavage, and sensitivity to large toxic molecules in *Escherichia coli*. J Bacteriol 184: 6093–6099.

36. Miller J.H (1992) A short course in bacterial genetics. A laboratory manual and handbook for Escherichia coli and related bacteria. CSHL Press.

37. Baba T, Ara T, Hasegawa M, Takai Y, Okumura Y, Baba M, et al (2006) Construction of *Escherichia coli* K-12 in-frame, single-gene knockout mutants: the Keio collection. Mol Syst Biol 2: 2006–008.

38. Sharan SK, Thomason LC, Kuznetsov SG, Court DL (2009) Recombineering: A homologous recombination-based method of genetic engineering. Nat Protoc 4: 206–223.

39. Uzzau S, Figueroa-Bossi N, Rubino S, Bossi L (2001) Epitope tagging of chromosomal genes in *Salmonella*. Proc Natl Acad Sci USA 98: 15264–15269.

