## Supplementary Information for "Glycan strand cleavage by a lytic transglycosylase, MltD contributes to the expansion of peptidoglycan in *Escherichia coli*"

Short title: Role of MltD in peptidoglycan expansion of *E. coli*

**S1 Text:**

**Supporting Materials and Methods:**

**Plasmid constructions:**

The genomic DNA from the WT strain, MG1655 was used as a template to amplify desired gene. PCR amplification was done using NEB Q5 high fidelity master mix. The clones that were obtained were confirmed by sequence analysis.

**pMK1:** This plasmid is a pTrc99a derivative carrying a cloned *mltD* gene under the *trc* promoter. *mltD* gene was cloned without its ribosome binding site using the following forward and reverse primers:

5'CGAGCTCATGAAGGCAAAAGCGATATTAC3' and

5'CGGGATCCTCAGGAATCTGGCATGTTGTTG3'. The resulting amplified DNA fragment was cloned using SacI-BamHI sites (underlined in the primer sequence) in a cloning vector pTrc99a to obtain pMK1.

**pMK2 and pMK3:** These are derivatives of pMK1 which have active site mutations in the *mltD* gene. For the generation of site directed variants of *mltD,* a three-step PCR was performed. For this, two primers that are complimentary to each other were synthesized with the desired mutation at the center. In the first PCR, N-terminal fragment of *mltD* gene was amplified using the forward primer used for pMK1 construction and the reverse primer containing the desired mismatch. In the second PCR, C-terminal fragment of *mltD* gene was amplified using forward mismatch containing primer and reverse primer that was used for pMK1 construction. The desired mismatches code for an alanine and lysine instead of a glutamate residue at position 125 (E125).

Primer sequence for making E125A is as follows (The changed codons are indicated in bold):

**pMK2 (E125A)**. 5’ GGTACTACTACCCATAGTG**GCG**AGCGCTTTTGATCCTCACG 3’ and 5’ CGTGAGGATCAAAAGCGCT**CGC**CACTATGGGTAGTAGTACC 3’

Sequence for making E125K is as follows:

**pMK3 (E125K).**

5’ AACTGGTACTACTACCCATAGTG**AAA**AGCGCTTTTGATCCTCACGCAAC 3’ and 5’ GTTGCGTGAGGATCAAAAGCGC**TTT**TCACTATGGGTAGTAGTACCAGTTC 3’

The first round of PCR was done using the common forward and SDM reverse and SDM forward and common reverse primer for each of the mutants to obtain two fragments. In the third step, these fragments were mixed in 1:1 molar ratio and used as template for another round of PCR using the common forward and reverse primers. The final PCR fragment was digested with SacI and BamHI and cloned into pTrc99a digested with the same enzymes. The plasmids pMK2 and pMK3 were confirmed for the mutations by sequencing.

**pMK4:** This is a variant of pMK1 carrying *mltD* gene without LysM domains. Truncated *mltD*$^{1-340}$ without *lysM* domains was cloned in pTrc99a. The forward primer was same as the one used for constructing pMK1 and the reverse primer sequence is 5’GGGATCCTCAGCTGTTAAGCGGCGTATTGTC 3’ containing the BamHI site. The PCR fragment was digested with SacI and BamHI and cloned in pTrc99a vector digested with the same enzymes. The truncation was confirmed by sequencing.

**pMK5:** *mltD* (without its promoter and RBS) having a C-terminal 1XFlag tag was constructed using the common forward primer used for construction of pMK1 and the reverse primer is of the following sequence:

5’CGGGATCCTTACTTGTCATCGTCATCCTTGTAATCGGAATCTGGCATGTTGTTG 3’

containing BamHI site. The PCR fragment was digested with SacI and BamHI and cloned in pTrc99a vector digested with the same enzymes. The fusion was confirmed by sequencing.

**pMK6:** The *mltD*$^{18-452}$ fragment lacking the N-terminal signal sequence (His-MltD) was cloned into pET28b vector using NdeI-BamHI sites. The forward and reverse primer sequences are as follows: 5’ GGAATTCCATATGAGTACCGGCAACGTTCAACAG 3’ and 5’ CGGGATCCTCAGGAATCTGGCATGTTGTTG 3’.

**Strain constructions:**

**Construction of deletion alleles**: Deletion strains used in the study were sourced from the Keio collection [1]. The presence of gene deletion was validated by PCR followed by sequence analysis. Deletions were introduced into the desired strains by P1 transductions. Wherever required, the Kan gene was flipped out using pCP20 plasmid.

**Construction of a strain carrying *mltD* gene under IPTG inducible promoter at its native chromosomal locus:** To engineer the cellular level of MltD, a strain was constructed in which the native promoter of *mltD* was replaced with a *lac* promoter which can be modulated by addition of IPTG. For the construction of this strain, hybrid primers with regions homologous to the chromosomal locus and a cassette containing a Cm$^{R}$- Lac promoter region of pHYD6012 were made. The regions homologous to chromosome are underlined: 5’GGTTAAGGTCAAAGAAAGATAGGTTCTGATAAAACTTTCTTGTGTAGGCTGGAGCTGCTTC 3’ and 5’CGAGCAGGACAGAGGCGAGTAATATCGCTTTTGCCTTCATTTCCTGTGTGAAATTGTTATCC 3’.

These primers were used to amplify the *lac* promoter sequence along with Cm$^{R}$ cassette from the plasmid pHYD6012 (a generous gift from Dr. Abhijit Sardesai, Centre for DNA Fingerprinting and Diagnostics, Hyderabad). The PCR fragment generated was electroporated into DY378 (a recombineering strain) [2] and colonies were obtained at 30°C on LB plates containing 15 µg/ml Cm. The construct was confirmed by PCR and sequencing. Finally, the construct was introduced into MG1655 through P1 transduction.

**Construction of *mltD*-Flag, *mltA*-Flag and *mltF*-Flag fusions:** The 3xFlag epitope tagging of *mltD*, *mltA* or *mltF* genes was done at the 3′ end at their native chromosomal locus as previously described [3]. Hybrid primers with regions homologous to the chromosomal loci and Flag-Kan cassette of pSUB11 were used. The regions homologous to chromosome are underlined and the region homologous to pSUB11 are italicized. The primer sequences are as follows:

***mltD*-3x Flag:**

5’AAGCTGACGTTGTTTGTGAAAAACAACAACATGCCAGATTCC*GACTACAAAGA CCATGACGG* 3’ and

5’AAGGCACCGGGGGAATCGGTGCCTTTTTATTATCTGGTTTG*CATATGAATATC CTCCTTAG* 3’

***mltA*-3x Flag:**

5’GTGCTGAAAACCGCCCCGGGCGCAGGTAACGTCTTTAGCGGC*GACTACAAA GACCATGACCGG* 3’ and

5’TCACCCTGTCATATCCGTAAAAACGGCATACAGAATATCACACA*CATATGAATA TCCTCCTTAG* 3’

***mltF*-3x Flag:**

5’TCTCTGCTGTTTTCCAGGAAAGGGAGTGAAGAGAAACAAAAT*GACTACAAAGA*

*CCATGACGG* 3’ and

5’<u>CCAGGAAATTAAAGCGCAGAAAAAGCGCAATCCTCGACGGA</u>*CATATGAATAT CCTCCTTAG* 3’

These primers were used to amplify the 3xFlag sequence from the plasmid pSUB11 with a $Kan^R$ marker flanked by FRT (flippase recognition target) sites. The PCR products were electroporated into DY378, and colonies were obtained at 30°C on LB plates containing 25 µg/ml Kan. The gene-Flag-$Kan^R$ region was transferred from DY378 into MG1655 by P1 transduction. These constructs were confirmed by PCR amplification, sequencing and western blotting using Flag antibodies. When required, the $Kan^R$ cassette was flipped out using pCP20, and the Kan-sensitive ($Kan^S$) derivatives of *mltD*-3xFlag were made.

**Construction of *mltD*::*lacZY*-Kan:** The *mltD*::Kan deletion of the Keio collection was converted into *lacZY*-Kan fusion as described earlier [4]. Initially, the deletion mutation was transferred into MC4100 strain (a strain carrying deletion of complete Lac operon) by P1 transduction. Next, the plasmid pCP20, encoding Flp recombinase was introduced into this strain to flip out the $Kan^R$ determinant at 30°C. Subsequently, into the $Kan^S$ derivatives carrying pCP20, an R6K based plasmid, pKGE137, encoding a promoter-less lacZY cassette along with $Kan^R$ marker flanked by FRT sites was introduced. $Kan^R$ colonies were selected on plates supplemented with X-Gal, and the presence of *lacZ*-Kan cassette at *mltD* locus was confirmed by PCR and sequencing. The *mltD*::*lacZY*-Kan fusion was then transferred into MC4100 or its *ΔrpoS* deletion derivative.

**Supplementary methods:**

### Protein overexpression

Overexpression plasmids encoding NlpI, Prc, MepS, MepM were described earlier [5,6]. MltD lacking signal sequence was constructed in this study. Overexpression of proteins was done using T7 RNA polymerase-based system as described earlier [5,6]. Overexpression of His-MltD was done in BL21 (λDE3) and a single transformant was grown overnight, and subcultured (1:100 ratio) into fresh LB broth containing respective antibiotics and grown till an $OD_{600}$ of 0.6 with shaking at 37ºC. Expression of MltD was induced by adding 50 µM IPTG (optimally standardized to achieve the highest amount of soluble protein) and grown further for 2 h at 37ºC. Cells were centrifuged and the pellet was then processed for purification of the expressed protein.

For the purification of His-MepS and Prc-His, NlpI-His, the plasmids were transformed into BL21 (λDE3) Δ*nlpI* strain and processed as described above with minor modifications:

His-MepS: Induced with 200 µM IPTG at 37ºC for 3 h.

MepM-His: Induced with 100 µM IPTG at 37ºC for 3 h.

Prc-His: Induced with 500 µM IPTG at 25ºC for 4 h.

NlpI-His: Induced with 500 IPTG at 37ºC for 3 h.

### Protein purifications

MepS, MepM, NlpI and Prc were purified as described earlier [5,6]. For MltD, the harvested cells were resuspended in 20 ml of lysis buffer (50 mM Tris pH8.0, 300 mM NaCl, 10 mM imidazole) and further lysed by sonication for 15-20 min (20%

Amplitude; 5-sec on-off cycle). The overexpressed proteins were soluble and did not require any detergent for solubilization. Cell debris was then removed by centrifugation at 30,000X RCF for 20 min at 4ºC and the supernatant was mixed with 0.5 ml of Ni2+- NTA agarose (Qiagen) at 4ºC for 1 h. This mixture was loaded into empty glass columns (Bio-Rad) and the supernatant was then allowed to pass through to retain the Ni-NTA beads inside the column. The beads were then washed with 30 ml of wash buffer-I (50 mM Tris pH 8.0, 300 mM NaCl, 30 mM imidazole, 1% Triton-X-100), then with 30 ml of wash buffer-II (50 mM Tris pH 8.0, 300 mM NaCl, 50 mM imidazole). Bound proteins were eluted with 15 ml of elution buffer (50 mM Tris pH 8.0, 300 mM NaCl, 250 mM imidazole). The eluted protein was buffer exchanged using a 30 kDa cut-off centrifugal membrane filter (Millipore) with 2X storage buffer (100 mM Tris pH8.0, 200 mM NaCl, 2 mM DTT) and concentrated. An equal volume of 100% glycerol (molecular grade) was added and protein sample was then stored at -30ºC. A small aliquot was run on SDS-PAGE to evaluate the purity of the protein.

**Pull down assays.** Pull downs (or co-purifications) were performed as described earlier [6]. Strains were grown overnight and next day, diluted 1:100 into 150 ml of LB and allowed to grow until $OD_{600}$ of 0.8. Cells were recovered by centrifugation, and the pellet was resuspended in 20 ml of lysis buffer (50 mM of Tris-Cl, 100 mM NaCl, 20% glycerol, 1% Triton X 100, 250 µg/ml lysozyme, 1X protease inhibitor, 20 units of DNase, 20 µg/ml of RNase and 10 mM imidazole; pH 8.0) and incubated in ice for 2 h. The mixture was sonicated, and the lysate was stirred overnight with glass beads at 4°C to solubilize the membrane proteins. Next day, the insoluble material of the lysate was removed by centrifugation at 15000 x g for 30 min. 100 µl from the supernatant was set aside as input fraction. To the rest of the supernatant,

300 µl of Ni-NTA agarose beads were added and mixed for 2 h at 4°C. The agarose beads were washed twice with wash buffer I (50 mM Tris-Cl, 100 mM NaCl, 1% Triton X 100, 20% glycerol and 20 mM imidazole; pH 8.0) and II (50 mM Tris-Cl, 100 mM NaCl, 20% glycerol and 50 mM imidazole; pH 8.0). The bound proteins were eluted using 250 µl elution buffer I (50 mM Tris-Cl, 100 mM NaCl, and 150 mM imidazole; pH 8.0) and II (50 mM Tris-Cl, 100 mM NaCl, 300 mM imidazole; pH 8.0). The input samples along with the elution fractions were subjected to SDS-PAGE and the proteins were detected by western blotting.

**β-galactosidase assay**

The assays were performed as described previously [7] using MC4100 *mltD::lacZ* strain or its derivatives. For measuring the beta-galactosidase values in stationary-phase cultures, the cell lysates were made as follows: Strains were grown overnight in LB medium, and next day diluted 1:100 into fresh LB medium and grown till $O.D_{600}$ of 3.0. 5 ml of the culture was taken, sonicated to lyse the cells and the lysate was then centrifuged at 14,000 RPM for 30 min. The supernatant was collected, and protein was estimated using Bradford reagent. Lysate equivalent of 10 µg of protein was taken and β-gal assay was performed using ONPG as the substrate as described earlier [7].

**Table 1. List of strains used in the study**

| Strain | Genotype | Source/ Reference |
|---|---|---|
| MG1655 | *rph*1 *ilvG rfb*-50 | Lab collection |
| DH5α | F⁻ *hsdR17 deoR recA1 endA1 phoA supE44 thi-1 gyrA96 relA1* Δ*(lac-argF)U169 φ80dlacZ* Δ*M15* | Lab collection |
| BW25113 | *lacI*$^{q}$ *rrnB3* Δ*lacZ4787* Δ(*araBAD*)567 Δ(*rhaBAD*)568 *hsdR*514 | [1] |
| DY378 | W3110 *λC1857* Δ(cro-bioA) | [2] |
| MC4100 | F− *araD139* Δ*argF-lac169* λ− e14−*flhD5301 relA1 rpsL150 rbsR22 deoC1* | Lab collection |
| BL21 (λDE3) | *ompT rB- mB-* (*PlacUV*5*::T*7-1) | Novagen |
| MR510 | Δ*mepS* Δ*mepM*/ pMN83 | [5] |
| MR508 | Δ*mepS* Δ*mepM* Δ*mepH*/ pMN83 | [5] |
| MR810 | *mepS*::*frt* | [5] |
| MK01 | Δ*mepS* Δ*mepM::Cm* | This study |
| MK02 | MK01 *prc** | This study |
| MK03 | MK01 *nlpI** | This study |
| MK04 | MK01 Δ*prc::*Kan | This study |
| MK05 | MK01 Δ*nlpI::*Kan | This study |
| MK06 | Δ*mepS* Δ*mepM* Δ*nlpI* (Kan$^{S}$)/ pMN83 | Lab collection |
| MK07 | MK06 Δ*mepA*::Kan | This study |
| MK08 | MK06 Δ*mepH*::Kan | This study |
| MK09 | MK06 Δ*dacB*::Kan | This study |

| | | |
|---|---|---|
| MK10 | MK06 Δ*pbpG*::Kan | This study |
| MK11 | MK06 Δ*ampH*::Kan | This study |
| MK12 | MK06 *mltA*::Kan | This study |
| MK13 | MK06 *mltB*::Kan | This study |
| MK14 | MK06 *mltC*::Kan | This study |
| MK015 | MK06 *mltD*::Kan | This study |
| MK016 | MK06 *mltE*::Kan | This study |
| MK017 | MK06 *mltF*::Kan | This study |
| MK018 | MK06 *mltG*::Kan | This study |
| MK019 | MK06 *digH*::Kan | This study |
| MK020 | MK06 *slt*::Kan | This study |
| MK022 | MR508 *prc*::Kan | This study |
| MK023 | MR508 *nlpI*::Kan | This study |
| MK024 | MK04 *mltD*::Kan | This study |
| MK025 | *mltD*-Flag-frt | This study |
| MK026 | Δ*nlpI*::*frt* MK025 | This study |
| MK027 | Δ*prc*::*frt* MK025 | This study |
| MK028 | Δ*nlpI* Δ*prc::frt* MK025 | This study |
| MK038 | *mepM::frt*/ pTrc99a | This study |
| MK039 | *mepM::frt*/ pMK1 | This study |
| MK040 | MG1655 $P_{lac}$::*mltD* | This study |
| MK041 | MR810 $P_{lac}$::*mltD* | This study |
| MK042 | *mltD*::*frt* | This study |
| MK043 | MR810 *mltD*::*frt* | This study |
| MK044 | MR510 *mltD*::Kan | This study |

| | | |
|---|---|---|
| MK045 | MG1655 Δ*lysA*::Kan | This study |
| MK046 | MR810 Δ*lysA*::Kan | This study |
| MK047 | MK042 Δ*lysA*::Kan | This study |
| MK048 | MK043 Δ*lysA*::Kan | This study |
| MK049 | *mepM*::*frt* Δ*lysA*::Kan | This study |
| MK050 | MK049 *mltD*::*frt* Δ*lysA*::Kan | This study |
| MK051 | *mltA*-3xFlag-Kan | This study |
| MK052 | MK051 Δ*nlpI*::*frt* | This study |
| MK053 | MK051 Δ*prc*::*frt* | This study |
| MK054 | *mltF*-Flag-Kan | This study |
| MK055 | MK054 Δ*nlpI*::*frt* | This study |
| MK056 | MK054 Δ*prc*::*frt* | This study |
| MK057 | MK025 Δ*rpoS*::Kan | This study |
| MK058 | MK026 Δ*rpoS*::Kan | This study |
| MK059 | MK027 Δ*rpoS*::Kan | This study |
| MK060 | MK025/pTrc99a | This study |
| MK061 | MK058/ pMK5 | This study |
| MK062 | MK026/pMK5 | This study |
| MK063 | MK026 Δ*rpoS*::Kan/pMK5 | This study |
| MK064 | MK027/pMK5 | This study |
| MK065 | MK027 Δ*rpoS*::Kan/pMK5 | This study |
| MK066 | MC4100 *mltD::LacZ* | This study |
| MK067 | MC4100 Δ*rpoS mltD::LacZ* | This study |
| MK068 | MK025 *degP*::Kan | This study |
| MK069 | MK025 *degQ*::Kan | This study |

| | | |
|---|---|---|
| MK070 | MK025 *ptrA*::Kan | This study |
| MK071 | MK026 *nlpI*::*frt* *prc*-HA-Cm/ pMN218 | This study |
| MK072 | MK025 *prc*-HA-Cm *mepS*::*frt*/ pMN217 | This study |
| MK073 | MK025 *prc*-HA-Cm *mepS*::*frt* *nlpI*::*frt*/ pMN217 | This study |
| MK083 | Δ*mepS* Δ*mepM::*Cm Δ*prc* | This study |
| MK084 | Δ*mepS* Δ*mepM::*Cm Δ*prc mltD::Kan* | This study |

**Table 2. List of plasmids used in the study**

| Plasmids | Relevant Features | Source/ Reference |
|---|---|---|
| pCA24N | $Cm^R$, *lacI*$^q$,$P_{T5\text{-}lac}$ | [8] |
| pCA24N-*mltA* | $Cm^R$, *lacI*$^q$,$P_{T5\text{-}lac}$::*mltA* | [8] |
| pCA24N-*mltC* | $Cm^R$, *lacI*$^q$,$P_{T5\text{-}lac}$::*mltC* | [8] |
| pCA24N-*mltD* | $Cm^R$, *lacI*$^q$,$P_{T5\text{-}lac}$::*mltD* | [8] |
| pCA24N-*mltE* | $Cm^R$, *lacI*$^q$,$P_{T5\text{-}lac}$::*mltE* | [8] |
| pCA24N-*mltF* | $Cm^R$, *lacI*$^q$,$P_{T5\text{-}lac}$::*mltF* | [8] |
| pCA24N-*mltG* | $Cm^R$, *lacI*$^q$,$P_{T5\text{-}lac}$::*mltG* | [8] |
| pCA24N-*digH* | $Cm^R$, *lacI*$^q$,$P_{T5\text{-}lac}$::*digH* | [8] |
| pCA24N-*slt* | $Cm^R$, *lacI*$^q$,$P_{T5\text{-}lac}$::s*lt* | [8] |
| pTrc99a | ColE1, $Amp^R$, *lacIq*, T7*lac* | Lab collection |
| pCP20 | pSC101(Ts), $Amp^R$, $Cm^R$, Flp | Lab collection |
| pMN83 | pBAD33-*mepS* | [5] |
| pET28b | T7 *lac* promoter; $Kan^R$ | Lab collection |
| pMN219 | pET21b-*prc*$^{23\text{-}682}$-His | [6] |
| pMN208 | pET21b-*nlpI*$^{19\text{-}294}$-His | [6] |
| pMN218 | pBAD18-*nlpI*-His, $Amp^R$ | [6] |
| pMN217 | pBAD18-*mepS*-His, $Amp^R$ | [6] |
| pMK1 | pTrc99a-*mltD* | This study |
| pMK2 | pTrc99a-*mltD*$_{E125A}$ | This study |
| pMK3 | pTrc99a-*mltD*$_{E125K}$ | This study |
| pMK4 | pTrc99a-*mltD*$^{1\text{-}340}$ | This study |
| pMK5 | pTrc99a-*mltD*-Flag | This study |
| pMK6 | pET28b-6X His-*mltD*$^{18\text{-}452}$ | This study |

**Supplementary Figures:**

Fig. S1

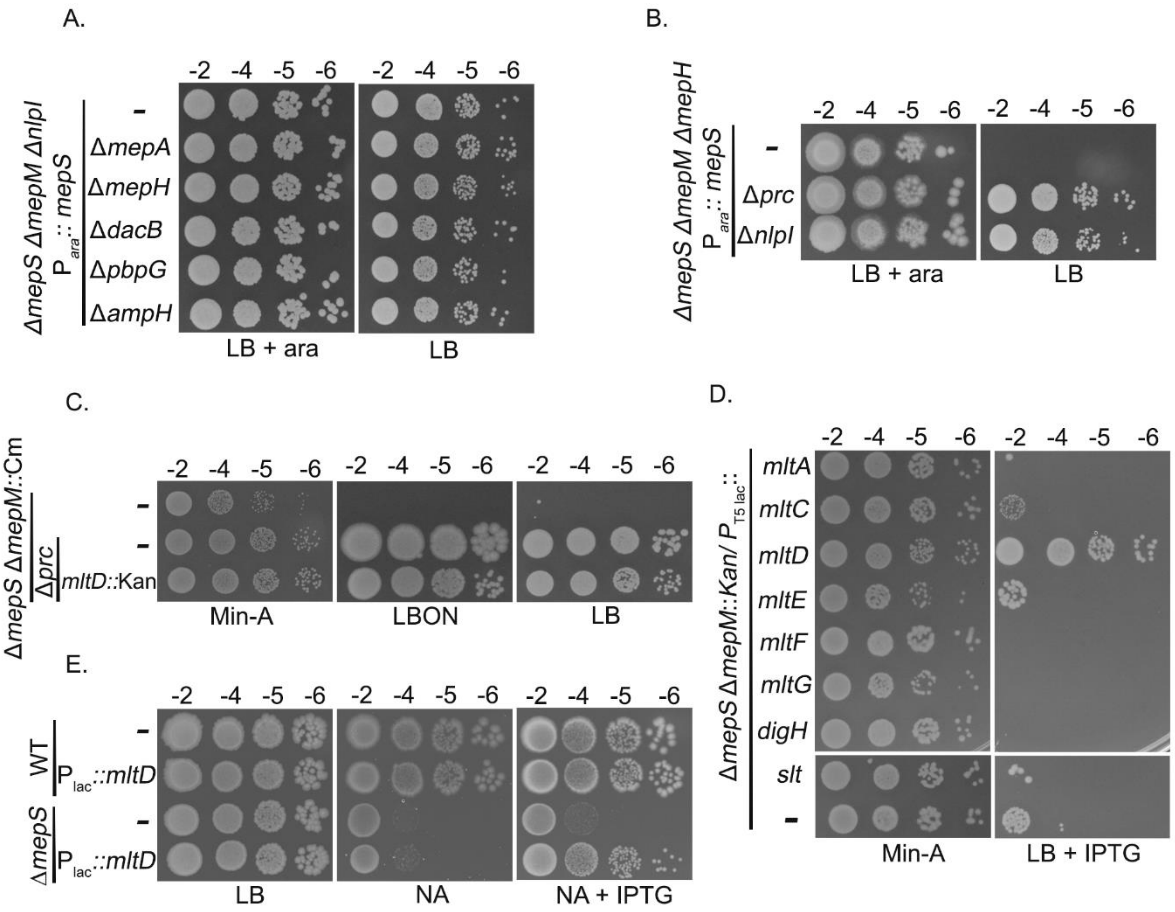


**Fig. S1. Absence of NlpI-Prc system rescues the growth defects of *mepS mepM* mutant through MltD.** (A) Δ*mepS* Δ*mepM* Δ*nlpI*/ P$_{ara::}$*mepS* and its derivatives carrying deletion of each of the known D,D-endopeptidases were tested for viability on LB with (0.2% ara) and without arabinose. Cells were grown overnight in LB with arabinose, serially diluted and 4 µl of each dilution was spotted on indicated plates and grown overnight. (B) Δ*mepS* Δ*mepM* Δ*mepH*/ P$_{ara::}$*mepS* and its mutant derivatives lacking *nlpI* or *prc* were tested for viability on LB with (0.2%) and without arabinose. (C) Δ*mepS mepM*::Cm double mutant or its derivatives were grown overnight in Min-A and their viability was tested on Min-A, LBON (LB without NaCl) and LB plates. (D) Growth of indicated strains was checked on Min-A and LB with IPTG (20 µM). (E) Cells of WT or *mepS* mutant having *mltD* under the control of *lac* promoter at its native chromosomal locus were grown overnight in LB and viability was checked on indicated plates with IPTG (10 µM).

Fig. S2

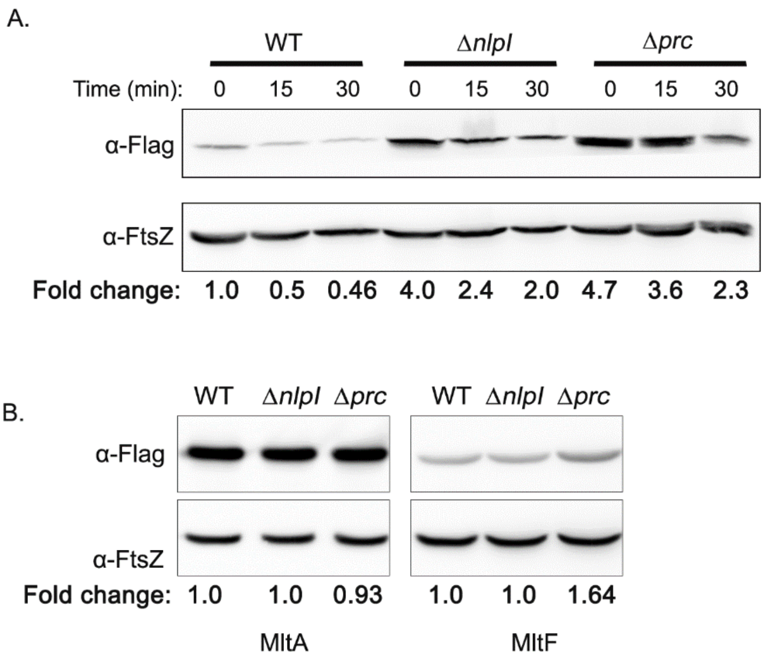


**Fig. S2. Regulation of LTs by NlpI-Prc proteolytic system.** (A) Determination of MltD half-life by spectinomycin-chase experiment. Indicated strains were grown in LB upto an $OD_{600}$ of 0.8-1.0 and 300 µg/ml spectinomycin was added to block translation. Fractions are collected at regular intervals, normalized to 1.0 OD, cell lysates were made, and aliquots equivalent of 0.5 OD were subjected to SDS-PAGE followed by western blotting. Bands were quantified using ImageJ software. FtsZ is used as a loading control to normalize the target protein. Fold change was calculated after normalization with FtsZ values. (B) WT and its mutant derivatives carrying *mltA*-Flag or *mltF*-Flag at their native chromosomal locus were grown in LB and fractions were collected between $OD_{600}$ of 0.8-1.0. The fractions were normalized to 1 OD, cell lysates were made and aliquots equivalent of 0.5 OD cells (for *mltA*-Flag) or 1 OD (in case of *mltF*-Flag) were subjected to SDS-PAGE followed by western blotting.

Fig. S3

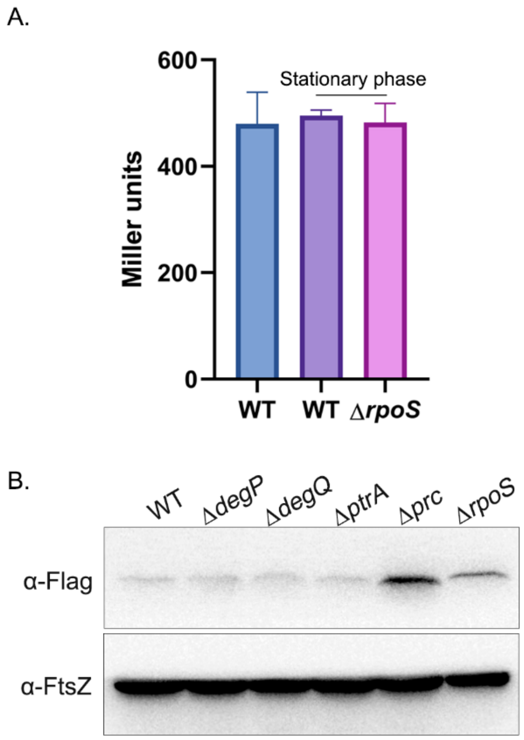


**Fig. S3. Post-transcriptional control of RpoS on MltD.** (A) β-galactosidase values of *mltD*::*lacZY* in WT and RpoS deletion mutant. Blue bar represents the β-gal values in log phase (at $OD_{600}$ of 0.6). The other two bars indicate the values of stationary-phase cells of WT and *rpoS* mutant. For this, the assay was done with cells grown to 3 OD. 5 ml of cells were collected, washed with PBS, sonicated and cell lysates were made. They were spun, supernatant was collected, and total protein content was estimated in each sample and equal amount of protein from each (10µg) lysate was used for β-galactosidase assay as described previously [7]. (B) Western blot showing expression of MltD-Flag in WT and its mutant derivatives as indicated. The experiment was carried out at 30°C as mutants of *degP* and *degQ* are sensitive to higher temperatures. Cell fractions were collected at 3 OD and normalized to 1.0 OD, cell lysates were made, and aliquots equivalent of 1 OD were subjected to SDS-PAGE and analysed by western blotting.

Fig.S4

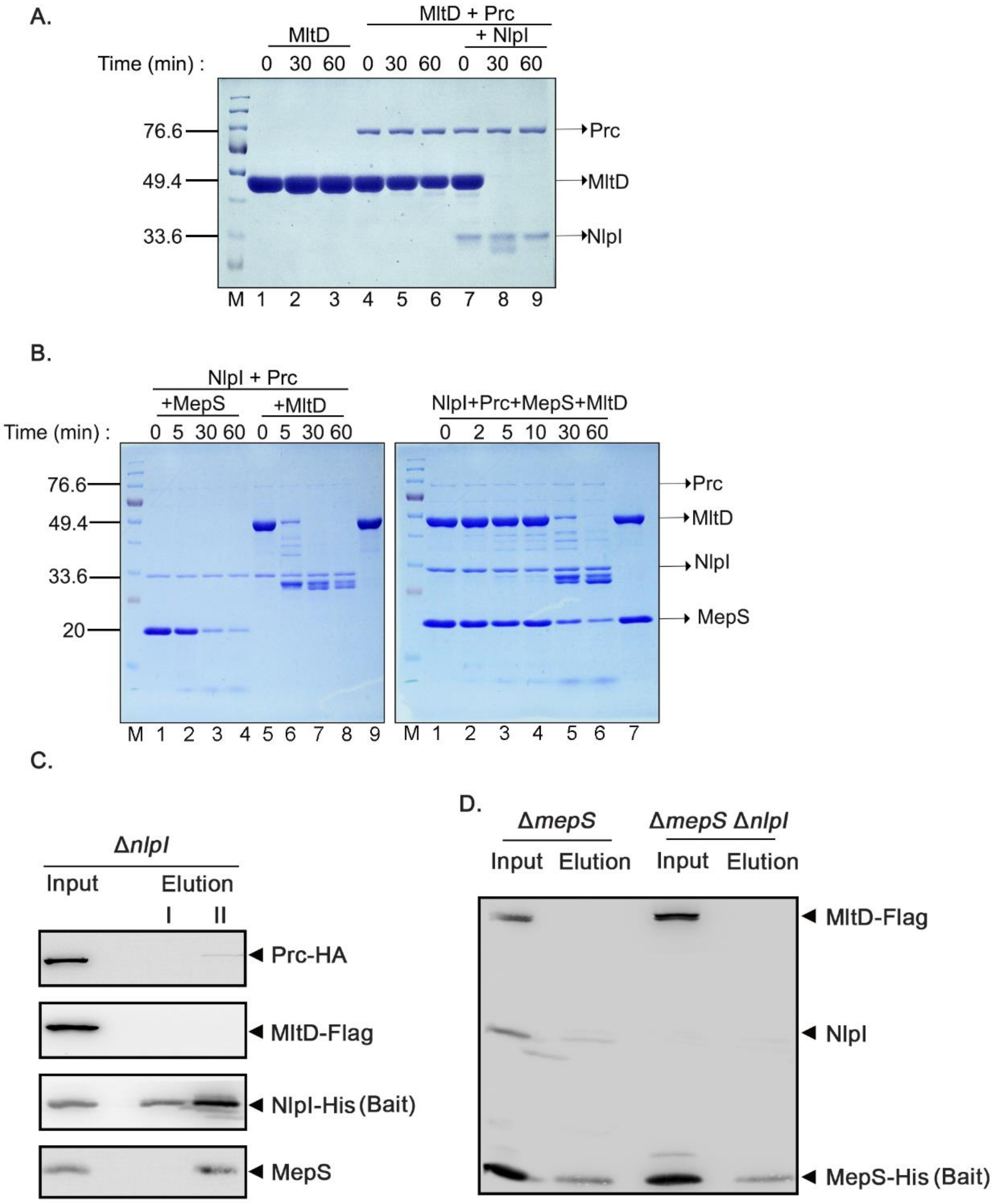


**Fig. S4. Interaction between MltD, NlpI and Prc proteins.** (A) *In vitro* degradation assay with purified MltD, Prc and NlpI proteins. Proteins were mixed in different combinations and incubated at 37°C followed by SDS-PAGE. Amounts of proteins used are as follows: MltD- 10 µg, NlpI- 1 µg, Prc- 0.4 µg. MltD protein itself was incubated to check for its stability. (B) *In vitro* degradation assay with purified MepS, MltD, Prc and NlpI proteins. Left panel shows degradation of MepS or MltD with NlpI and Prc proteins. Right panel shows the degradation of MepS and MltD co-incubated with NlpI and Prc. (C) *In vivo* pull-down assay to check the interaction of NlpI with MltD. Plasmid borne NlpI-His was used as a bait to examine its interaction with MltD-Flag. Prc-HA and MepS are used as positive controls. (D) *In vivo* pull-down assay with plasmid borne MepS-His was used as a bait to check its interaction with chromosomal MltD-Flag in WT and *nlpI* deletion mutant.

**Supplemental References:**


1. Baba T, Ara T, Hasegawa M, Takai Y, Okumura Y, Baba M, et al (2006) Construction of *Escherichia coli* K-12 in-frame, single-gene knockout mutants: the Keio collection. Mol Syst Biol 2: 2006-008.

2. Sharan SK, Thomason LC, Kuznetsov SG, Court DL. 2009. Recombineering: A homologous recombination-based method of genetic engineering. Nat Protoc 4: 206–223.

3. Uzzau S, Figueroa-Bossi N, Rubino S, Bossi L. 2001. Epitope tagging of chromosomal genes in *Salmonella*. Proc Natl Acad Sci USA 98 Proc Natl Acad Sci USA 98: 15264–15269.

4. Ellermeier CD, Janakiraman A, Slauch JM. 2002. Construction of targeted single copy lac fusions using λ Red and FLP-mediated site-specific recombination in bacteria. Gene 290: 153-161.

5. Singh SK, Saisree L, Amrutha RN, Reddy M (2012) Three redundant murein endopeptidases catalyse an essential cleavage step in peptidoglycan synthesis of *Escherichia coli* K12. Mol Microbiol 86:1036–1051.

6. Singh SK, Parveen S, SaiSree L, Reddy M (2015) Regulated proteolysis of a cross-link-specific peptidoglycan hydrolase contributes to bacterial morphogenesis. Proc Natl Acad Sci USA 112:10956–10961.S1

7. Miller J.H. 1992. A short course in bacterial genetics. A laboratory manual and handbook for *Escherichia coli* and related bacteria. CSHL Press.

8. Kitagawa M, Ara T, Arifuzzaman M, Loka-Nakamichi T, Inamoto E, Toyonaga H et al (2005) Complete set of ORF clones of *Escherichia coli* ASKA library (a complete set of *E. coli* K-12 ORF archive): unique resources for biological research. DNA Res. 12(5): 291–299.
